# Pangenome analytics reveal two-component systems as conserved targets in ESKAPEE pathogens

**DOI:** 10.1101/2020.04.25.061994

**Authors:** Akanksha Rajput, Yara Seif, Kumari Sonal Choudhary, Christopher Dalldorf, Saugat Poudel, Jonathan Monk, Bernhard O. Palsson

**Affiliations:** Systems Biology Research Group, Department of Bioengineering, University of California, San Diego, San Diego, CA, United States; Bioinformatics and Systems Biology Program, University of California, San Diego, San Diego, CA, United States; Department of Pediatrics, University of California, San Diego, San Diego, CA, United States; Novo Nordisk Foundation Center for Biosustainability, Technical University of Denmark, Kemitorvet, Building 220, 2800 Kongens, Lyngby, Denmark

**Keywords:** Two-component systems, ESKAPEE pathogens, antibiotic resistance, Pangenomic analysis, genomic architecture

## Abstract

Bacteria sense and respond to environmental stimuli through two-component systems (TCSs), that are composed of histidine kinase sensing and response regulator elements. TCSs are ubiquitous and participate in numerous cellular functions. TCSs across the ESKAPEE pathogens, representing the leading causes of nosocomial infections, were characterized using pangenome analytics, including annotation, mapping, pangenomic status, gene orientation, sequence variation, and structure. Our findings fall into two categories. 1) phylogenetic distribution of TCSs: (i) the number and types of TCSs varies between species of the ESKAPEE pathogens; (ii) TCSs are group-specific, i.e., Gram-positive and Gram-negative, except for KdpDE; (iii) most TCSs are conserved among genomes of an ESKAPEE, except in *Pseudomonas aeruginosa*. 2) sequence variation: (i) at the operon level, the genomic architecture of a TCS operon stratifies into a few discrete classes; and (ii) at the gene sequence level, histidine kinases, responsible for signal sensing, show sequence and structural variability as compared to response regulators that show a high degree of conservation. Taken together, this first comprehensive pangenomic assessment of TCSs reveals a range of strategies deployed by the ESKAPEE pathogens to manifest pathogenicity and antibiotic resistance. It further suggests that the conserved features of TCSs makes them an attractive group of potential targets with which to address antibiotic resistance.

## Introduction

Two-component systems (TCSs) are universally distributed among bacterial species (Mitrophanov and Groisman 2008; Gross, Aricò, and Rappuoli 1989). They participate in numerous cellular processes including signaling and pathogenicity (Zschiedrich, Keidel, and Szurmant 2016) and also play a major role in the pathogenicity of the highly infectious ESKAPEE group of pathogens, which is an acronym for *Enterococcus faecium, Staphylococcus aureus, Klebsiella pneumoniae, Acinetobacter baumannii, Pseudomonas aeruginosa, Enterobacter spp*. and *Escherichia coli* (Boucher et al. 2009; Pendleton, Gorman, and Gilmore 2013). The ESKAPEE pathogens, consisting of both Gram-positive and Gram-negative bacteria, are the leading cause of nosocomial life threatening infections and are in the WHO’s “priority pathogen” list (Santajit and Indrawattana 2016). The problem of trying to tackle nosocomial infection worsens due to the increase in antibiotic resistance and virulence.

The histidine kinase (HK) and response regulator (RR) are two important components of TCSs (Ibrahim, Puthiyaveetil, and Allen 2016). HK is a transmembrane protein which senses external signals (Bhagirath et al. 2019; West and Stock 2001). Upon sensing the environmental stimuli, the conserved histidine residue gets autophosphorylated by receiving gamma-phosphate from ATP. Further, the phosphate is transferred to aspartate residues of the response regulator (Schaller, Kieber, and Shiu 2008). Upon phosphorylation, the response regulator undergoes structural changes, which further helps in the expression of various target genes (Gao, Bouillet, and Stock 2019). Therefore, the changes in target gene expression mediates cellular expression to respond to the external stimuli (**Figure 1A**). Thus, TCSs help bacteria to acclimatize to a wide range of external factors.

**Figure 1.**
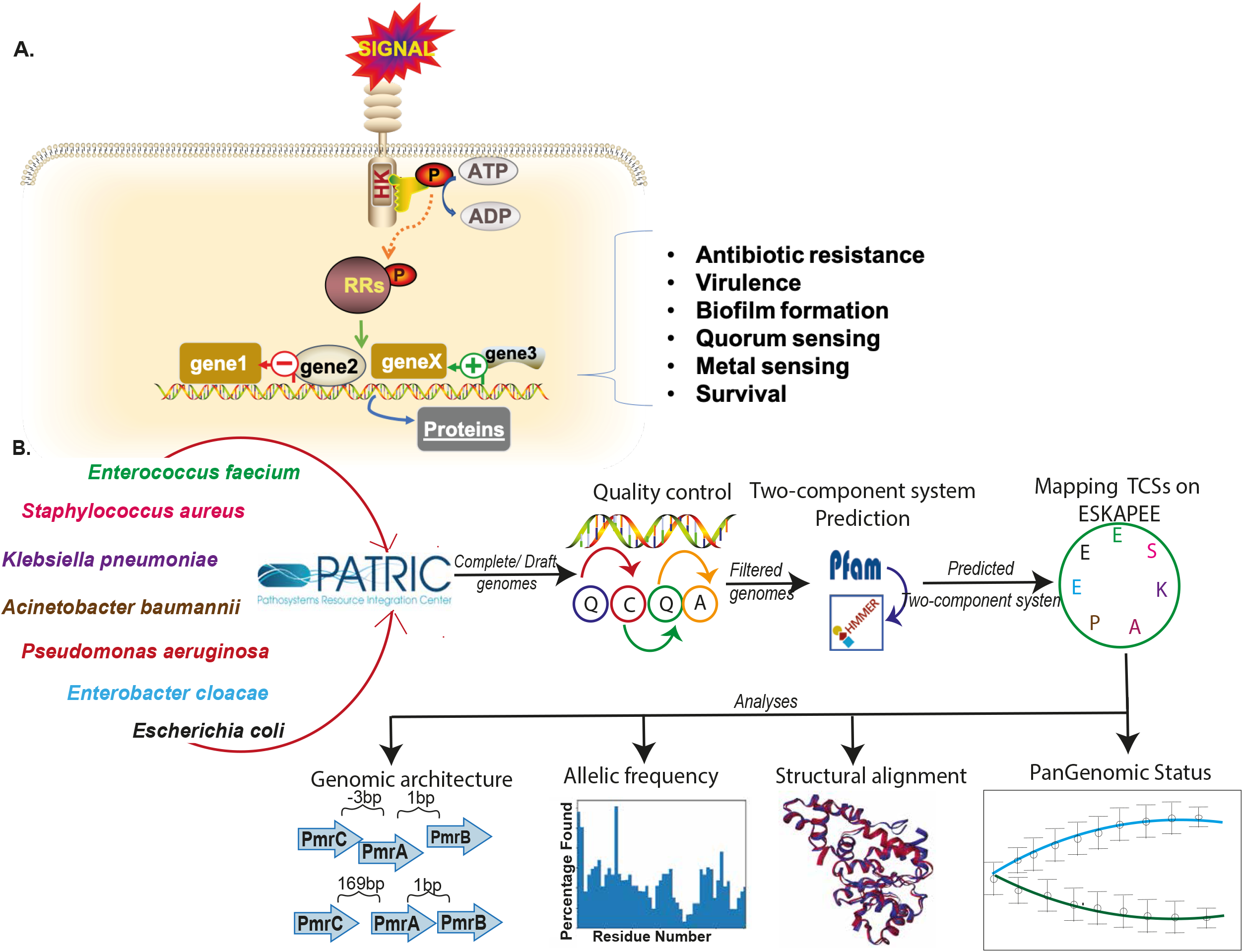
Pangenome analysis of two-component systems: A) Schematic diagram depicting the mechanism of a two-component system. B) Flow chart showing the methodology used in the study.

TCSs are involved in antibiotic resistance, virulence, quorum sensing, biofilm formation, metal sensing, motility, survival, and many other functions (Bhagirath et al. 2019; Rajput, Kaur, and Kumar 2016). The antibiotic resistance TCSs help bacteria to survive in the presence of various antibiotics (Tierney and Rather 2019). The TCSs involved in virulence help bacteria to sustain in the host or at the site of pathogenicity (Barrett and Hoch 1998). The quorum sensing, motility, and biofilm related TCSs allow bacteria to communicate, move, and form colonies to survive in unfavorable environments (Barrett and Hoch 1998; Prüß 2017). Further, bacteria also have TCSs in order to survive in other conditions like high pH, metals, anaerobic conditions, nutrient sensing, etc. (Bhagirath et al. 2019; Golby et al. 1999). Therefore, the many roles played by TCSs make them a valuable potential target for antimicrobials. Several studies have confirmed this potential (Reading et al. 2009; Kato and Groisman 2004).

Among all the functions of TCSs, antibiotic resistance has been the most extensively investigated in bacteria, especially within the ESKAPEE group of pathogens (Santajit and Indrawattana 2016)(Tierney and Rather 2019)(Santajit and Indrawattana 2016). Bacteria adapt different TCS mechanisms to express antibiotic resistance phenotypes (Muller, Plésiat, and Jeannot 2011). The mechanisms include over expression of efflux pumps, cell surface modifications, upregulation of antibiotic resistance genes, and increased biofilm formation (Tierney and Rather 2019; Cerqueira et al. 2014). Various strategies need to be developed to overcome these specialized modifications against antibiotics in bacteria.

TCSs are a fundamental determinant of bacterial physiological states. Despite being ubiquitous and vital for bacterial survival, TCSs have not yet been the subject of a detailed pangenomic analysis. A pangenomic study would be helpful to understand the conservation status of all the TCSs involved in antibiotic resistance, virulence, biofilm, motility, and others involved in the basic survival mechanisms in bacteria. Literature shows that TCSs could be a promising target to fight pathogenicity of bacteria, especially antibiotic resistance (Worthington, Blackledge, and Melander 2013). This pangenome study, driven by the availability of a large number of strain-specific genome sequences, is focused on exploring all TCSs among the ESKAPEE pathogens.

## Material and Methods

The overall methodology is provided in Figure **1B** and is described in detail below.

### Collection and quality control of ESKAPEE genomes

The ESKAPEE genomes were downloaded from the Pathosystems Resource Integration Center (PATRIC) *v*3.5.43 (Wattam et al. 2017). The downloaded genome has “Complete” and “Draft” genome status, “human, *Homo sapiens”* host, and “good” genome quality. Further, the five levels of quality control were done to get a more refined set of genomes for downstream analysis. First, the genomes annotated as “Plasmid” were removed. Second, the genomes that didn’t have Multilocus sequence typing (MLST) were removed. MLST filtration is important to have only the genomes with the presence of housekeeping genes to provide good resolution of genome characterization. Third, only those genomes with a number of contigs<100 were retained, to confer good quality assembly. Fourth, genomes with the coding region of genes i.e. CDS between [Average ± 2(Standard deviation)] were kept, to get rid of the mis-annotated genomes. Fifth, the genomes with a number of N’s > 1000 were filtered out. The table depicts the resulting ESKAPEE pathogen genomes at each quality control step is provided in **Supplementary Figure S1,2,3**.

### Annotation of two-component systems among the ESKAPEEs

The Hidden Markov Model (HMM) (Eddy 1996) and BLAST (Altschul et al. 1997) were used to annotate the TCSs among all the ESKAPEE pathogens. The HMM profile information of HK and RR were collected from MIST3.0 (Gumerov et al. 2020), P2CS (Ortet et al. 2015), and literature. The Pfam profiles of the RR and HK in all ESKAPEE pathogens were downloaded using the Pfam32.0 (El-Gebali et al. 2019). The Pfam profiles are the summarized output of protein sequences of the family and built through seed and automatically generated full alignment (Finn et al. 2008). Further, these HMM profiles were used to annotate the TCSs among all ESKAPEE using hmmsearch tool.

### Summarizing the two-component systems among the ESKAPEEs

The annotated TCSs of ESKAPEE were curated and summarized. The summarization of TCSs were done broadly in four categories i.e. Antibiotic resistance, Virulence, Others/general, and Predicted/Unknown function. All the TCSs were scanned for their frequency of occurrence among the individual pathogens. Afterward, four heatmaps were constructed for above-mentioned categories with the information of the frequency of occurrence of the TCSs among them.

### Pangenomic analysis of two-component systems among the ESKAPEEs

We performed a pan genomic analysis of all the TCSs by checking their distribution among strains. Further, the frequency distribution of the TCSs in all or atleast 98% strains considered as core, some strains (accessory), or only one strain (unique) (Monk et al. 2013). The distribution was calculated as (Strain with the presence of TCSs/Overall strains)*100.

For each species, we plotted proxy pan and core genome curves as in (Seif et al. 2018) but limiting our input to TCSs. Briefly, we generated 1,000 random permutations of the input genomes, and for each permutation, we randomly sampled strains one at a time without replacement. At the first draw, we counted the number of TCSs detected. At the next draw, we counted the number of TCSs, but subdivided them into three counts: 1) the core count: the number of unique TCSs found in both draws; 2) the pan count: the total number of unique TCSs when pooling the two draws, and; 3) the new TCSs count: the number of TCSs found in the second draw that we couldn’t find in the first draw. This process was repeated until all strains were drawn. We generated a vector of recorded set sizes for each of the 1,000 permutations, and calculated the average and standard deviation for each step. We then fit Heap’s law (an empirical power law) to the vector of new gene sets, and calculated the mean and standard deviation of the fitted parameters α and k. Heaps’ law was originally developed to describe the count of unique words in a text as a function of the length of the text. Here, it can be expressed as *n* = *k N*^−α^, where n is the total number of genomes, N is the total count of new TCSs discovered at each draw, k is a multiplicative constant and α is the gene discovery decay rate (Tettelin et al. 2008). The pan genome can be described as either “closed” (α > 1) or “open” (α < 1). A pan genome is “open” when the pan count increases indefinitely as new genomes are considered, and “closed” when the rate of increase of the pan count slows down as more strains are analyzed and the pan count eventually reaches a plateau (at which point, no new genes are discovered).

### Sequence variation among two-component systems among the ESKAPEEs

The sequences for the RR and HK of TCSs were used for the analysis. Further, the BLASTp (Altschul et al. 1997) was run between the sequences and the respective reference sequence. Any insertions, deletions, or SNP’S between the RR or HK sequences and the reference sequence were counted as a variant residue at the residue number of the reference sequence.

The sequence variation among the RR and HK sequences were also done using the Principal Component Analysis (PCA) plots. The important peptide features like amino acid composition, dipeptide composition, and tripeptide composition were calculated (Rajput, Gupta, and Kumar 2015). Further, these features were used to make PCA plots for RR and HK in all ESKAPEE pathogens.

### Structural alignment of two-component systems among the ESKAPEEs

The protein structures of HK and RR of the two-component system involved in antibiotic resistance (VraSR) among *E. faecium* and *S. aureus* species were used for structure alignment. The PDB (Wang 2012) doesn’t have the protein structure for most of the HK and RR components for ESKAPEE two-component systems. Thus, we opted for VraSR. The protein structures were downloaded from the PDB database (*e.g*. 4GT8, 4GVP, and 5HEV). However, the VraS protein structure for *E. faecium* was constructed using I-TASSER from sequence (UniProt ID: S4DWF2). Further, the VraS and VraR were aligned using FATCAT (Ye and Godzik 2004). The structure alignment resulted in the form of Root mean square deviation (RMSD). The lower RMSD value between aligned structures represents the more structural similarity between the two, while more RMSD refers to more variability among structures.

### Genomic architecture of two-component systems among the ESKAPEEs

The genomic architecture provides an important idea about the spatial arrangement of the genes in an operon (Choudhary et al. 2018). Here we constructed the genomic architecture of the most shared and important TCSs among the categories like antibiotic resistance, virulence, and others/general. For example, PmrAB, VraSR, BaeSR (Antibiotic resistance); AgrCA, WalKR, AlgZR (Virulence); CusSR, KdpDE (Others (general)) TCSs. The genome architecture was constructed after scanning the genomes of pathogens, TCSs operon genes, calculation of intergenic distances, and orientations. All this information was collated and depicted in the form of arrow diagrams.

## Results

### Annotation of two-component systems

Different numbers of TCSs were annotated among ESKAPEE pathogens using the HMM approach (**Figure 2B**). We categorized the TCSs into four different groups, namely antibiotic resistance, virulence, others (general), and predicted family. The categorization was done as per the major function of the TCSs reported in literature. However, the “predicted family” includes the TCSs whose family has been annotated rather than the exact TCS. The detailed list of TCSs and their functions among ESKAPEE is provided in **Supplementary Tables S1-6**.

**Figure 2.**
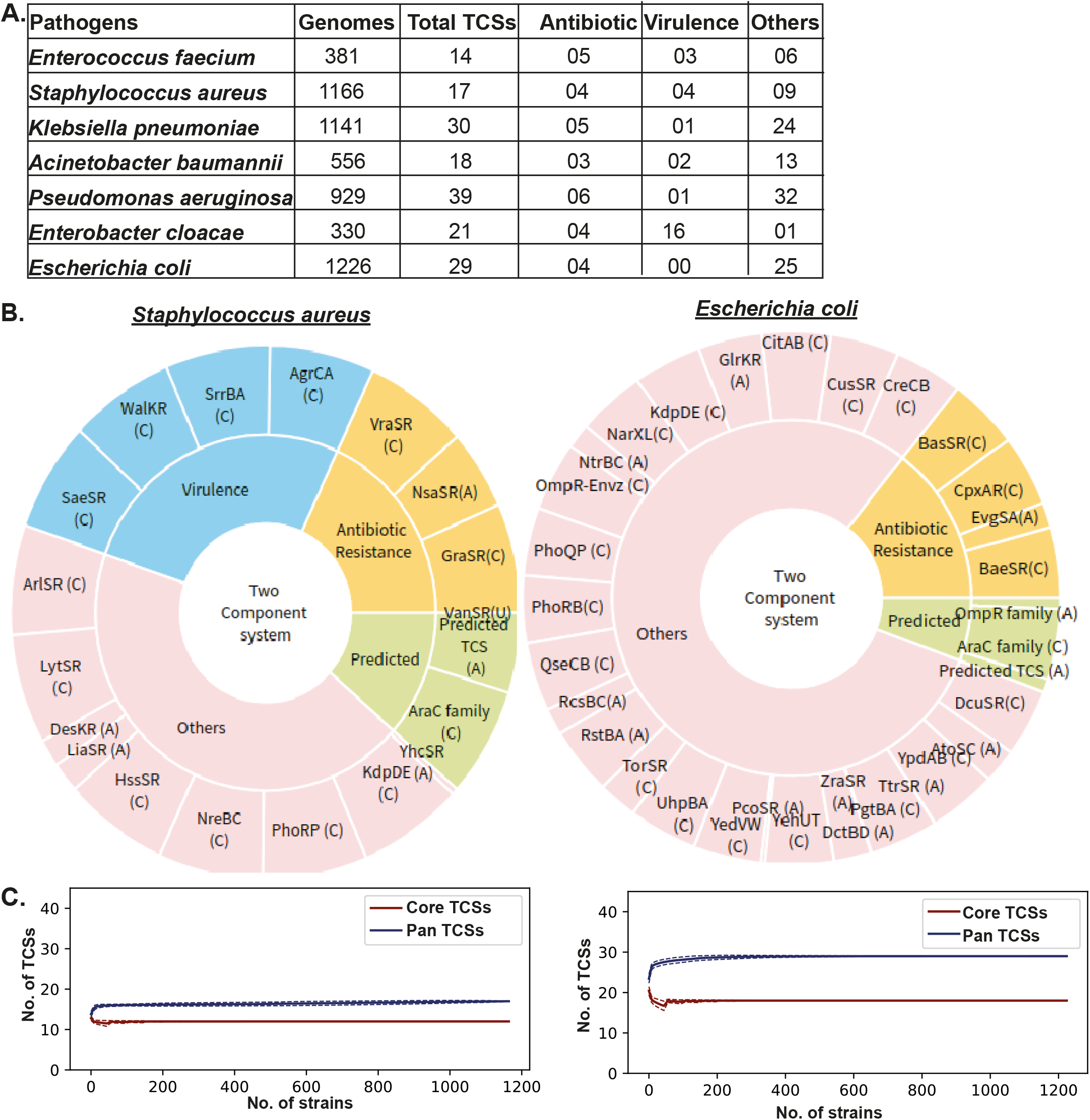
Pan genome analysis of two-component systems. A) Table showing the total number of genomes after quality control and two-component systems annotated and categorised in antibiotic resistance, virulence, and others. B) Multilevel pie chart depicting the distribution of TCSs in all four categories in S. aureus, and E. coli. C) Pangenome curves for S. aureus and E. coli. The curve shows the conservation status of core and pan-genome for TCSs.

The highest number of TCSs (39) were mapped in *P. aeruginosa*, with 6 functioning as antibiotic resistance, 1 for virulence, with the remaining 32 falling into the others (general) category. Among ESKAPEE pathogens, *E. faecium* possesses the lowest number of TCSs (14), with 5 as antibiotic resistance. Other ESKAPEE pathogens like *K. pneumoniae* (30), *E. coli* (29), *E. cloacae* (21), *A. baumannii* (18), and *S. aureus* (17) mapped with different TCSs (**Figure 2A**). The highest number of TCSs involved in antibiotic resistance are present in *P. aeruginosa*, while the highest number of TCSs for virulence are found in *E. cloacae*. The TCSs with other (general) functions are most abundant in *P. aeruginosa*.

### Pangenome analysis of two-component systems

The pangenome analysis of the TCSs among the ESKAPEE pathogens showed that most of the TCSs are part of the “core” genome, i.e., they are shared across the genome (**Figures. 2B and S4**). Apart from their distribution as core, the TCSs are also found as an “accessory” component. We found only two TCSs that were “unique” to a genome: VanSR in *S. aureus* and CprSR in *P. aeruginosa*. The conservation status of the TCSs is also depicted as the pangenome curve showing core and pangenome TCSs (**Figures. 2C and S5**).

Our first goal was to characterize the level of conservation of the two-component systems across species. We constructed core and pan-genome curves focused on the TCSs for each species (Methods). Briefly, the core genome curve corresponds to the number of conserved TCSs, and the pan genome curve reflects the total number of TCSs as more strains are taken into account. This is the first attempt to categorize TCSs into the core and the pan genome. Our initial categorization is focused on five criteria.

1) The number of TCSs found in core genomes of ESKAPEE pathogens: We find that the number of TCSs which are part of the core genome (i.e., present in more than 98% of genomes of a species, see methods) varies across species. In total, *P. aeruginosa* strains have the largest number of core TCSs (*n* = 21), followed by *E. coli* (n = 18), *K. pneumoniae* (n = 17), *S. aureus* (n = 12), *A. baumannii* (n = 6) and *E. faecium* (n = 5). Surprisingly, none of the TCSs are part of the *E. cloacae* core genome (**Figure. 3A**).

**Figure 3.**
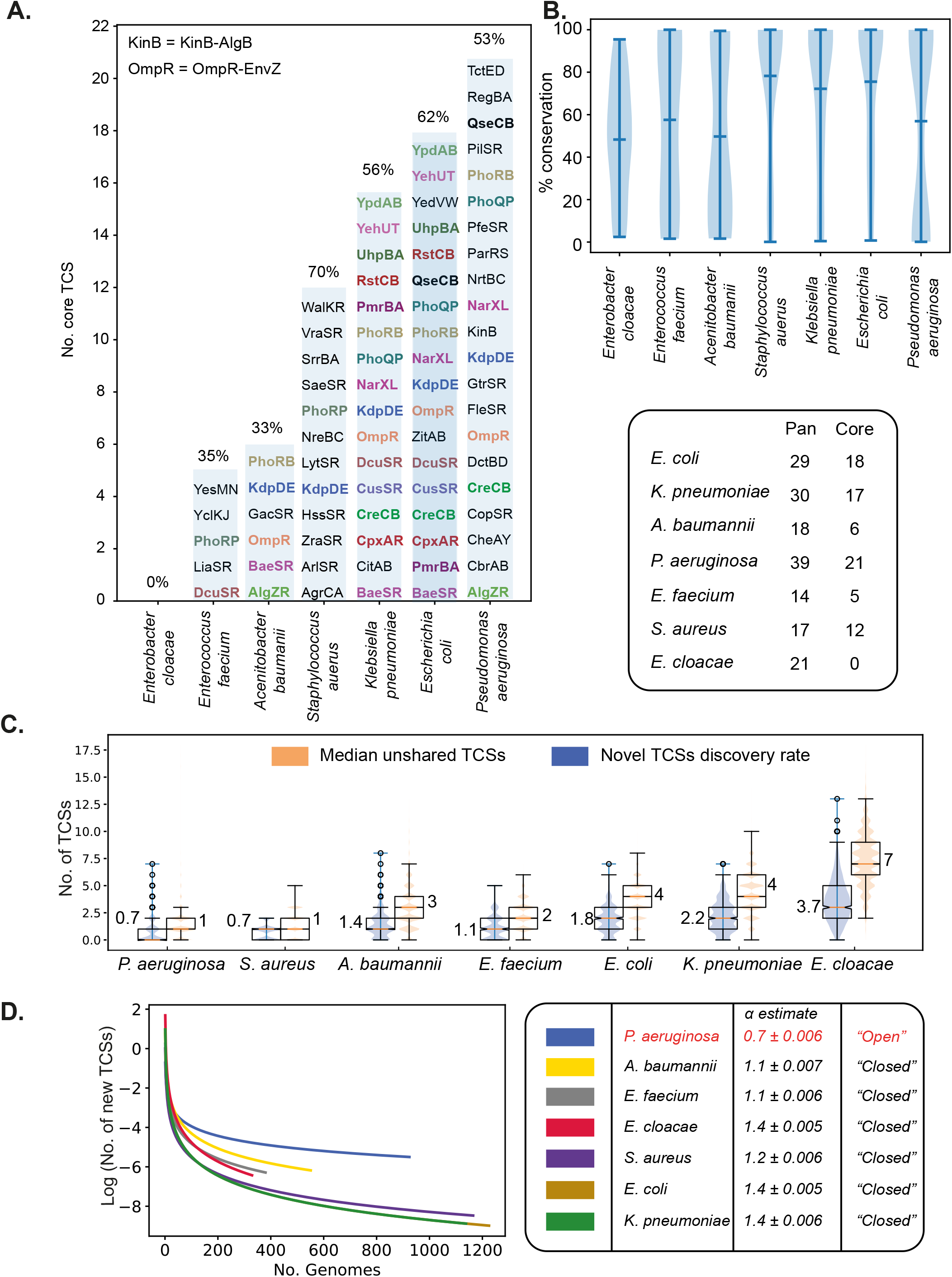
Pan genome analysis of two-component systems: A) Core TCSs across species. A core TCS is defined as a two-component system gene present in more than 98% of the strains. The percentage of TCSs that are part of the core is displayed on top of each bar. The common TCSs are shown with same colors. B) TCSs are variably conserved across strains. The percentage of strains in which a TCS is present is calculated for each TCS, and the distribution of percentages is plotted for each species. C) TCS discovery curves. The number of new TCSs discovered as more strains are taken into consideration decreases across species. Heap’s law was fitted to each curve, and the decay rate was estimated. A decay rate that is larger than 1 indicates a closed pan genome. P. aeruginosa is the only species with a decay rate smaller than 1, suggesting that the number of TCSs are unbounded, and that new genes will constantly be discovered as new P. aeruginosa genomes are sequenced. In contrast, the set of TCSs in all six other species is bounded and ceases to increase as more strains are sequenced. D) Median unshared TCSs and novel gene discovery rate at step one of the gene discovery curves in C. The novel TCS discovery rate represents the average number of new two-component systems discovered when two strains are drawn randomly, and the gene content of the second strain is compared to that of the first strain. The median unshared TCSs represents the number of two-components that differ between two strains (i.e. the difference between the intersection and the union of the two sets).

2) Common TCSs among ESKAPEE pathogens: The TCSs were mapped and depicted in the form of heatmaps to summarize their shared and unshared status along with pangenomic status among ESKAPEE pathogens. The summary of TCSs involved in antibiotic resistance and virulence are provided in **Figure. 4A** with predicted family, and others (general) in **Figure. S6**. Most of the TCSs are shared among the pathogens. For example, the antibiotic resistance TCS, PmrBA, is shared among *K. pneumoniae*, *A. baumannii*, *P. aeruginosa*, *E. cloacae*, and *E. coli*. The TCS involved in virulence, AlgZR, is found in *A. baumannii* and *P. aeruginosa*. The KdpDE TCSs, which is involved in others (general) functions, is distributed among *S. aureus*, *K. pneumoniae*, *P. aeruginosa*, *E. cloacae*, and *E. coli* (**Figure. 4A**). However, the function of certain core TCSs are similar across species.

**Figure 4.**
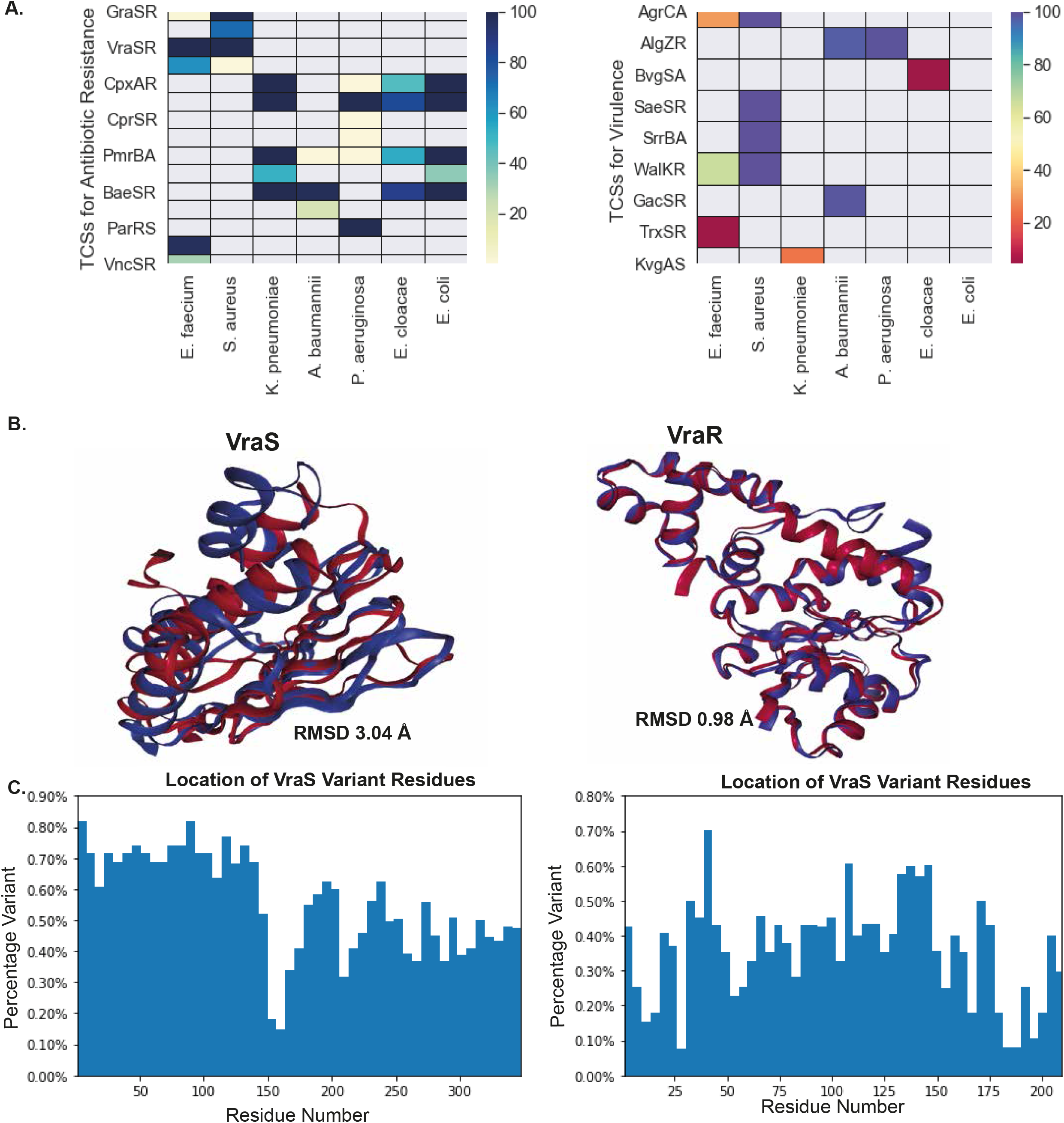
Pan genome analysis of two-component systems. A) Heatmaps depicting the TCSs involved in Antibiotic resistance, Virulence, and Other (general). The color of boxes is in accordance with the distribution of TCSs in the strains of respective ESKAPEE pathogens. B) Structural alignment of the Histidine kinase and Response regulator for VraSR two component system. The Root Mean Square Deviation (RMSD) of the aligned structure shows the similarity among them. If the RMSD is 0 it means the aligned structures are similar, while a high value of RMSD means dissimlarity in structures. C) Sequence variants bar graphs of VraS and VraR TCSs. The graph is plotted between percentage variation versus Number of residues.

3) Percentage of TCSs found in the core genomes of given ESKAPEE pathogens: While *P. aeruginosa* has the largest number of core TCSs, the proportion of core TCSs versus pan TCSs is highest in *S. aureus* (70%). In fact, the percentage of strains sharing any one of the TCSs varies greatly within and across species, with a generally high percentage of conservation in *S. aureus* (78%), *K. pneumoniae* (72%) and *E. coli* (75%) (**Figure. 3B**). In contrast, a TCS is shared only in 48%, 58%, and 50% of strains, on average, in *E. cloacae, E. faecium*, and *A. baumannii*, respectively. The distribution of percent conservation of TCSs is bimodal in *P. aeruginosa*.

4) Pangenomic status of TCSs for a given ESKAPEE: We investigated whether the set of TCSs was finite across a species, and whether we would continue to discover new TCSs as new strains are sequenced. For this purpose, we fitted Heaps’ law to a curve plotting the number of new genes discovered as more strains are taken into account (**Figure. 3C**, Methods). Two parameters, α and k, are estimated when fitting Heaps’ law. When α < 1, we consider the pan genome to be “open”, i.e., we would expect to find new TCSs as more strains are sequenced indefinitely. This condition only applied to the new gene discovery curve of *P. aeruginosa*, revealing that the set of TCSs is finite in all of the other species.

5) TCSs are shared between the two strains of the same species: We plotted the average number of new TCSs discovered when a second strain is examined, and the number of unshared genes between any two strains (**Figure. 3D**). Despite having the largest α, *P. aeruginosa* strains had the lowest average number of unshared TCS genes (n = 1), and the lowest new TCSs discovery rate (0.7), while *E. cloacae* had the highest values in both the number of unshared TCSs (n = 7) and novel TCS discovery rate (3.7).

### Genomic architecture of two-component systems

We scanned the genomic architecture of the most frequently shared TCSs among ESKAPEEs with antibiotic resistance, virulence, and others (general) categories and found that it varies (**Figure. 5** and **Figures. S8,9**). However, we also found some variation in gene arrangement within the same bacterial strains e.g., PmrBA, WalKR, and KdpDE TCSs, as shown in **Figure. 5**. Upon comparing the variation in gene arrangement in the TCS operon within each category, we found that more variation exists among TCSs in the others (general) category as compared to those involved in virulence and antibiotic resistance.

**Figure 5.**
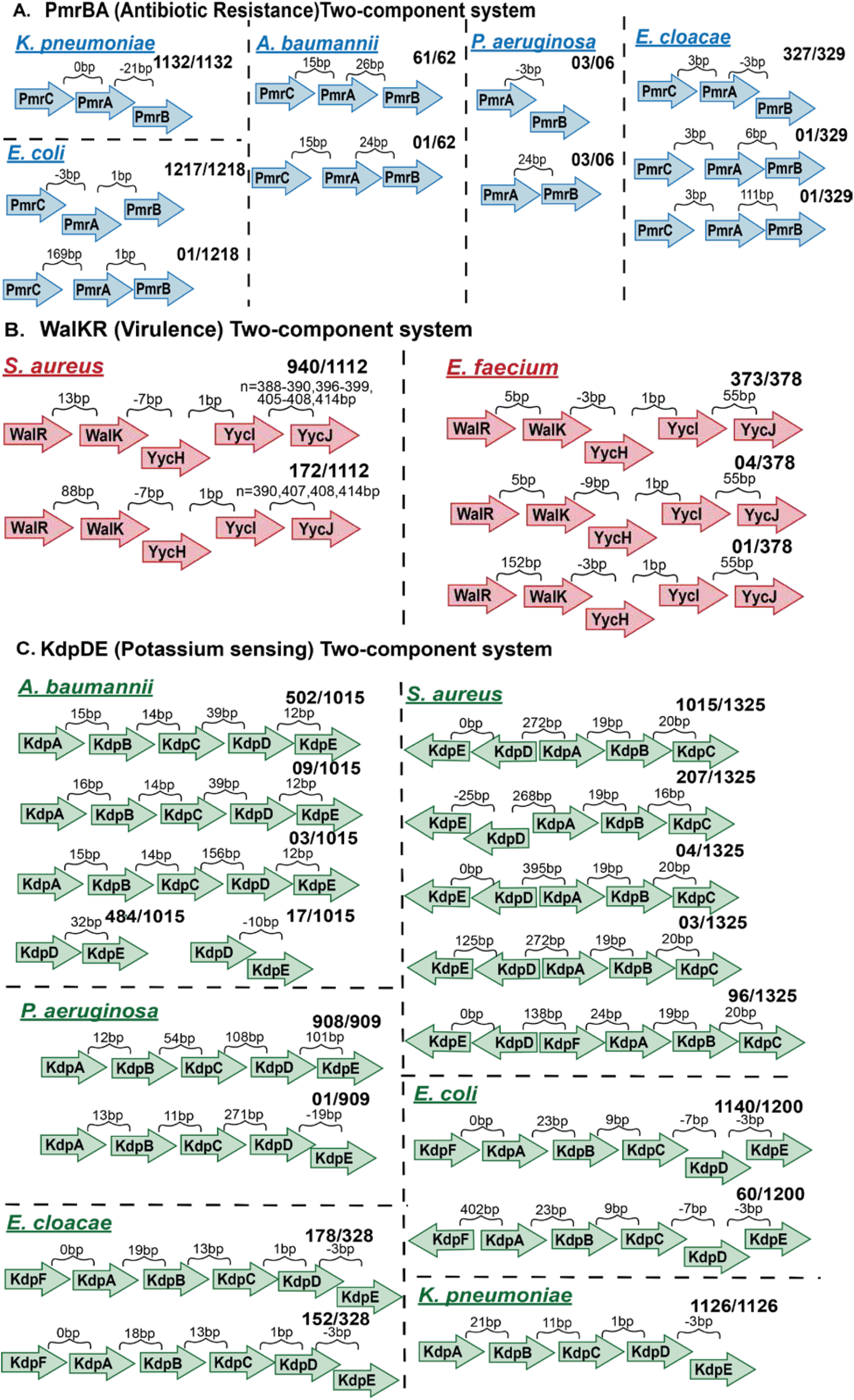
Pan genome analysis of the two-component systems shows a discrete number of classes. The direction of arrows in the TCS operon genes is the representation of those present in the positive strand. A similar arrangement is present in the negative strand. The length of the arrows is a representation of genes, not the scale. A) Genomic architecture of PmrBA two-component system involved in antibiotic resistance among Gram-negative ESKAPEE pathogens: K. pneumoniae, A. baumannii, P. aeruginosa, E. cloacae, and E. coli. B) Genomic architecture of the WalKR two-component system involved in virulence. The WalKR system is found in Gram-positive ESKAPEE pathogens: E. faecium, and S. aureus. C) Genomic architecture of KdpDE potassium (K^+^) sensing two-component system in S. aureus, K. pneumoniae, A. baumannii, P. aeruginosa, E. cloacae, and E. coli.

For example, the PmrBA two-component system has three genes in the operon: PmrB, PmrA, and PmrC. The PmrBA is found in five gram-negative ESKAPEE pathogens: *E. coli*, *E. cloacae*, *P. aeruginosa*, *K. pneumoniae*, and *A. baumannii*. The PmrBA operon shows different intergenic distances in these five pathogens despite them performing the same antibiotic resistance function. Likewise, the intergenic distances and gene arrangement varies among the bacteria in the WalKR and KdpDE two-component systems. For example, the WalKR operon is found in *E. faecium* and *S. aureus* while KdpDE is in *S. aureus, K. pneumoniae, A. baumannii, P. aeruginosa, E. cloacae*, and *E. coli*.

### Sequence and structural variation among the two-component systems

The sequence and structural variations were checked in histidine kinases and response regulator components of the TCSs. The sequences of both the components were checked to discover the percentage variation among them (**Figures. 4C and S7A**). For VraSR, VraS (HK) and VraR (RR) have variant scores of 0.27 and 0.18. In WalKR, the variant score in WalK (HK) and WalR (RR) is 0.12 and 0.05, respectively. In general, the HK domain shows more variation as compared to RR. Among the HK domain, the C-terminus shows more variability than the C-terminus. Additionally, the sequence variation of RR and HK among ESKAPEE pathogens was checked and depicted in the form of 3D PCA plots. For example, the 3D PCA plots of *S. aureus* and *A. baumannii* are depicted in **Figure. S4B**. The RR sequences of the respective TCSs seem to be tightly clustered as compared to HK. Taken together, the sequence variation analysis reflects that HK has more sequence variation as compared to RR in the ESKAPEE pathogens.

We also looked at the structural variation among HK and RR domains of TCSs. The structural alignment of VraS and VraR is provided in **Figure. 4B**. The structure alignment resulted in the form of root mean square deviation (RMSD). The lower RMSD value between aligned structures represents more structural similarity between the two, while higher RMSD is indicative of more variability among structures. The VraR alignment of *S. aureus* and *E. faecium* showed a RMSD of 0.98 Å. However, the VraS protein alignment of *S. aureus* and *E. faecium* showed a RMSD of 3.04 Å. HK has a higher RMSD value as compared to RR, thus HK showed more structural variability in structure than to RR.

## Discussion

In this study, we carried out a pan-genome analysis of TCSs in ESKAPEE pathogens. The study was made possible due to the recent growth in the number of strain-specific sequences available for these pathogens. With respect to phylogenetic distribution of TCSs, we find that the number of TCSs varies among ESKAPEE pathogens and they are group specific, i.e., among Gram-positive and Gram-negative, except in the case of KdpDE. Most TCSs are conserved among the pathogens (found in the closed pan genome) except in the case of *P. aeruginosa*. With respect to sequence and structural variation, we find that TCS operons are stratified in discrete classes, which is more pronounced in TCSs involved in general functions. The histidine kinases that sense environmental signals show more variability as compared to response regulators, which maintain cellular expression.

The ESKAPEE pathogens possess different categories of TCSs (see Tables S1-S7). The number and types of TCSs reflects the characteristics of the particular bacterium. For example, most of the TCSs in *P. aeruginosa* are related to biofilm formation while in *A. baumannii* they deal with metal sensing. We found that the majority of TCSs are shared among the two major bacterial groups (Gram-positive or Gram-negative bacteria), while fewer of them are exclusive to an individual ESKAPEE pathogen (Bourret and Silversmith 2010; Barrett and Hoch 1998). Pangenomic analysis of TCSs allows us to decipher their phylogenetic distribution and conservation.

The TCS pangenomes of most ESKAPEEs are found to be closed, which adds to their value as potential conserved targets for a species (Barrett et al. 1998). Pangenome analysis further shows that various TCSs are common to more than one ESKAPEE pathogen, including: VraSR (Antibiotic resistance); AlgZR (Virulence); and CitAB, PhoRP, and UhpBA (others (general)). Thus, these TCSs could serve as candidates for broad-spectrum inhibitors (Worthington, Blackledge, and Melander 2013). However, some TCSs were also part of the variant, or accessory, pangenome, which is present in a particular subset of strains.

The closed ESKAPEE TCS pangenomes reflect their conservation status and should make them good targets with regard to pathogenicity and antibiotic resistance. *P. aeruginosa* has the highest number of TCSs in the core component of the pangenome. Surprisingly, *P. aeruginosa* strain CLJ1 seems to be an outlier, because it carries a total of 33 TCSs, five of which are unique to this strain (including BfmSR, CarSR, CprSR, MifSR, and RoxSR), and eight of which are shared across less than 10% of *P. aeruginosa* strains (including BfiSR, CpxAR, CzcSR, PirSR, PmrBA, PprAB, RcsCB, and RocS2A2). CLJ1 was isolated in 2010 from the lungs of a patient with fatal hemorrhagic pneumonia in France, and contains an elevated number of ISL3-family insertions affecting major virulence-associated phenotypes and increased antibiotic resistance (Sentausa et al. 2019).

While the shared TCSs among different bacterial species exhibit the same function, the genomic architecture differs. The intergenic distances within the genes in an operon are thought to be evolutionarily conserved among a broad range of prokaryotes (Okuda et al. 2007). However, we found the genomic arrangement of the TCS operons fall into discrete classes. In a previous study, the *agr* operon in *S. aureus* was shown to fall into discrete classes that correlated with the host range of a given strain (Choudhary et al. 2018). In this study we show that the genomic architectures of TCS operons generally fall into discrete classes, which are more pronounced in the TCSs performing other (general) functions (**Figure. 3**).

Histidine kinases and response regulators comprise a TCS. The HK is membrane bound while the RR is its cytoplasmic counterpart (West and Stock 2001). HK genes are found to be more sequence variable than RR genes. Further, the structural alignment of both the domains among different ESKAPEE species further confirms more variation in HK components (see **Figure. 4**). The HK sequence and structural variation is especially pronounced in its N-terminal domain likely due to its function as a sensor for a broad range of environmental signals. Our results are in agreement with previous studies which show that the transmembrane N-terminals in HKs are responsible for signal sensing while the cytoplasmic C-terminal helps with phosphate transfer (Capra and Laub 2012).

As antibiotic resistance represents a major health concern worldwide, there is a growing need to identify new and promising targets in pathogenic bacteria. This first comprehensive pangenomic study of TCSs confirms their conservation and universality among ESKAPEE pathogens. Given that TCSs are integral mechanisms that enable antibiotic resistance, virulence, and basic metabolic functions, they could be targeted to tackle pathogenicity and reduce antibiotic resistance among nosocomial infections caused by ESKAPEE pathogens.

## Code Availability

The code used in the Analysis of the study is available at https://github.com/akanksha-r/TCS_Pangenome

## Acknowledgements

We thank Marc Abrams for reviewing the manuscript and providing constructive suggestions. This work was supported by NIH Grant U01 AI124316 and Novo Nordisk Foundation Grant NNF10CC1016517.

## Author contributions

A.R. and B.O.P. designed research; A.R., Y.S., and K.S.C. performed research; A.R., Y.S., K.S.C., C.D., S.P., and J.M. performed analyses. A.R., and Y.S. wrote the manuscript. All the authors have read and approved the manuscript.

## Notes

### Competing Interest Statement

The authors have declared no competing interest.

