## Supplementary for "Pangenome analytics reveal two-component systems as conserved targets in ESKAPEE pathogens"

**Supplementary information**

**S1. Supplementary Tables**

**Supplementary Table S1:** Detailed list of Two-component systems and their functions among *Enterococcus faecium*

**Supplementary Table S2:** Detailed list of Two-component systems and their functions among *Staphylococcus aureus*

**Supplementary Table S3:** Detailed list of Two-component systems and their functions among *Klebsiella pneumoniae*

**Supplementary Table S4:** Detailed list of Two-component systems and their functions among *Acinetobacter baumannii*

**Supplementary Table S5:** Detailed list of Two-component systems and their functions among *Pseudomonas aeruginosa*

**Supplementary Table S6:** Detailed list of Two-component systems and their functions among *Enterobacter cloacae*

**Supplementary Table S7:** Detailed list of Two-component systems and their functions among *Escherichia coli*

**S2. Supplementary Figures**

***Supplementary Figure S1:*** *Flow charts depicting the quality control steps and scatter plots for A) Table showing total number of genomes filtered at each step quality control steps, B) Enterococcus faecium, B) Staphylococcus aureus*

***Supplementary Figure S2:*** *Flow charts depicting the quality control steps and scatter plots for A) Pseudomonas aeruginosa, B) Escherichia coli, C) Klebsiella pneumoniae*

***Supplementary Figure S3:*** *Flow charts depicting the quality control steps and scatterplots for A) Enterobacter cloacae, B) Acinetobacter baumannii*

***Supplementary Figure S4:*** *Pan genome analysis of the two-component systems in form of Multilevel pie chart depicting the distribution of TCS in all the four categories in Enterococcus faecium, Klebsiella pneumoniae, Acinetobacter baumannii, and Enterobacter cloacae, and Pseudomonas aeruginosa*

***Supplementary Figure S5:*** *Pan genome analysis of the two-component systems in form of the Pangenome curve of Enterococcus faecium, Klebsiella pneumoniae, Acinetobacter baumannii, and Enterobacter cloacae, and Pseudomonas aeruginosa. The curve shows the status of core and pan-genome for TCS. All the four pathogens show "closed" pangenome for TCS except P. aeruginosa*

***Supplementary Figure S6:*** *Pan genome analysis of the two-component systems: A) Heatmaps depicting the TCS involved in others (general) category, B) Heatmaps depicting the predicted family of TCS*

***Supplementary Figure S7:*** *Pan genome analysis of the two-component systems: A) Sequence variants bar graphs of WalK and WalR TCS. The graph is plotted between percentage variation v/s Number of residues, B) Principal component Analysis (PCA) curves showing histidine kinase and response regulators of Staphylococcus aureus and*

*Acinetobacter baumannii. The PCA curves plotted by using peptide features i.e. amino acid composition, dipeptide composition, and tripeptide composition*

***Supplementary Figure S8:*** *Pan genome analysis of the two-component systems: A) Genomic architecture of BaeSR antibiotic resistance two-component system among Gram-negative ESKAPEE pathogens i.e. K. pneumoniae, A. baumannii, E. cloacae, and E. coli. B) Genomic architecture of VraSR antibiotic resistance two-component system. The VraSR system found in Gram-positive ESKAPEE pathogens i.e. E. faecium, and S. aureus. C) Genomic architecture of AgrCA virulence two-component system in E. faecium, and S. aureus*

***Supplementary Figure S9:*** *Pan genome analysis of the two-component systems: A) Genomic architecture of AlgZR virulence two-component system among Gram-negative ESKAPEE pathogens i.e. A. baumannii, and P .aeruginosa. B) Genomic architecture of CusSR Copper sensing two-component system. The CusSR system found in K. pneumoniae, A. baumannii, E. cloacae, and E. coli*

### S1. Supplementary Tables

***Supplementary Table S1:*** *Detailed list of Two-component systems and their functions among Enterococcus faecium*

| **TCS (*E. faecium*)** | **Function** | **References** |
|---|---|---|
| CiaHR/YycGF/VickR | Daptomycin resistance, cell wall homeostasis | (Diaz et al. 2014; Tran et al. 2013; Panesso et al. 2015) |
| GraSR | Cationic antimicrobial peptides | (Diaz et al. 2014) |
| VanSR | Vancomycin resistance | (Arthur, Molinas, and Courvalin 1992; Hong et al. 2008) |
| VncSR | Vancomycin resistance | (L. Hancock and Perego 2002) |
| VraSR | Vancomycin resistance | (Belcheva and Golemi-Kotra 2008; Bem et al. 2015) |
| YesMN | Growth phase adaptation, carbohydrate utilization | (L. Hancock and Perego 2002) |
| DcuSR | Fumarate & succinate regulation | (Zientz, Bongaerts, and Unden 1998; Unden, Wörner, and Monzel 2016) |
| YclKJ | Unknown | (L. Hancock and Perego |

| | | |
|---|---|---|
| | | 2002) |
| PhoRP | Phosphate sensing | (Santos-Beneit 2015) |
| PhoRB | Stress response, virulence | (Bem et al. 2015) |
| LytSR | Autolysin, regulates extracellular murein hydrolase activity, bacterial cell death and pyruvate utilization | (L. Hancock and Perego 2002; Ali et al. 2017) |
| WalKR | Virulence, Cell wall homeostasis, Cell division | (Fakhruzzaman et al. 2015; Dubrac et al. 2008) |
| TrxSR | Virulence, Regulation of bacteriocin genes | (Armstrong et al. 2016; Leday et al. 2008) |
| AgrCA | Virulence, Exo- and cell wall protein synthesis | (L. Hancock and Perego 2002; L. E. Hancock and Perego 2004) |

***Supplementary Table S2:*** *Detailed list of Two-component systems and their functions among Staphylococcus aureus*

| **TCS (*S. aureus*)** | **Function** | **References** |
|---|---|---|
| GraSR | Cationic antimicrobial peptides resistance | (Yang et al. 2012; Muzamal et al. 2014) |
| NsaSR (BceSR/BrsSR) | Nisin sensitivity associated response, stress response, cell wall synthesis, membrane transport | (Mensa et al. 2014; Makarova et al. 2018) |
| VraSR | Vancomycin/cell wall antimicrobials sensing, Cell wall biosynthesis | (Yin, Daum, and Boyle-Vavra 2006; Kuroda et al. 2003) |
| VanSR | Vancomycin resistance | (Hong et al. 2008; Macielag et al. 1998) |
| ArlSR | regulates agglutination and autolysin | (Walker et al. 2013; Bronner, Monteil, and Prévost 2004) |
| LytSR | Antibiotic tolerance, biofilm formation, stationary phase survival, cell death and analysis | (Sharma-Kuinkel et al. 2009) |
| DesKR | Membrane Fluidity, regulated transcription | (J. W. Kim et al. 2016) |
| LiaSR | cell wall peptidoglycan biosynthesis, antibiotic, acid, and detergent stresses | (Shankar et al. 2015; Belcheva and Golemi-Kotra 2008) |
| HssSR | Iron sensing, intracellular heme homeostasis | (Stauff, Torres, and Skaar 2007; Torres et al. 2007) |
| NreBC | oxygen-responsive nitrogen regulation | (Villanueva et al. 2018; Schlag et al. 2008) |

| | | |
|---|---|---|
| PhoRP | Sense Phosphate, monitor cell wall teichoic acid metabolism, hemolysin expression | (Botella et al. 2014; Burnside et al. 2010) |
| KdpDE | Potassium sensing | (Xue et al. 2011; Zhao et al. 2010) |
| YhcSR/ AirSR | Oxygen sensing, regulates the lac and opuCABCD Operon | (J. Sun et al. 2005; Yan et al. 2012) |
| AgrCA | Exo- and cell wall protein synthesis, quorum sensing | (Choudhary et al. 2018; R. P. Novick et al. 1995) |
| SaeSR | Secreted factors mostly involved in immune evasion | (Giraudo et al. 1999; Richard P. Novick and Jiang 2003) |
| SrrBA | virulence, activation of genes involved in anaerobic metabolism, cytochrome and haem biosynthesis and down-regulation of agr-RNAIII | (Pragman et al. 2004; Kinkel et al. 2013; Christmas et al. 2019) |
| WalKR | Bacterial cell envelope composition, cell viability and controls major autolysin genes involved in cell wall degradation, biofilm formation | (Dubrac et al. 2007; Delauné et al. 2012) |

***Supplementary Table S3:*** *Detailed list of Two-component systems and their functions among Klebsiella pneumoniae*

| **TCS (*K. pneumoniae*)** | **Function** | **References** |
|---|---|---|
| KvgAS | Virulence | (C.-T. Lin and Peng 2006) |
| EvgSA | Multidrug-related, free radical stresses, sensing iron-limiting conditions | (Lai et al. 2003; Ramos et al. 2016) |
| BaeSR | Multidrug resistance, lipopolysaccharide modifications regulation | (Bhagirath et al. 2019) |
| CpxAR | Cefepime and chloramphenicol resistance, osmolarity regulation | (Srinivasan, Vaidyanathan, et al. 2012; Ramos et al. 2016) |
| PhoQP | Polymyxin resistance, L-aminoarabinose synthesis regulation | (S. Y. Kim, Choi, and Ko 2014; Jayol et al. 2015) |
| PmrBA | Colistin and polymyxin resistance, lipopolysaccharide modification | (S. Sun et al. 2009; S. Y. Kim, Choi, and Ko 2014) |
| KdpDE | Potassium transport mechanism | (Freeman, Dorus, and Waterfield 2013; Hirakawa, Nishino, Hirata, et al. 2003) |
| CusSR | Copper sensing and induces CusCFBARS efflux system, silver resistance | (Zahid, Zulfiqar, and Shakoori 2012; Hanczvikkel et al. 2018) |

| | | |
|---|---|---|
| CreCB | Carbon source responsive, ß-lactam resistance response | (Lautenbach et al. 2001) |
| AtoSC | Acetocetate sensing | (Kumar et al. 2011) |
| PhoRB | Stress response, virulence | (Srinivasan, Venkataramaiah, et al. 2012) |
| RstBA | Response to environmental stimuli | (S.-C. Chen et al. 2013) |
| GlrKR | Regulates Envelop homeostasis | (Szklarczyk et al. 2019) |
| DcuSR | Fumarate respiration | (Golby et al. 1999) |
| PcoSR | Copper resistance sensing | (Wu et al. 2019; Y.-T. Chen, Chang, Lai, et al. 2004) |
| UhpBA | Regulates hexose phosphate metabolism | (Verhamme et al. 2002) |
| QseCB | Quorum sensing, Motility | (Bem et al. 2015; Weigel and Demuth 2016) |
| NarXL | Nitrate/Nitrite reduction | (Bem et al. 2015) |
| GlnLG | Nitrogen regulation, Glutamate metabolism | (Verhamme et al. 2002; Lai et al. 2003; Ninfa and Magasanik 1986) |
| NifLA | Regulates Nitrogen fixation (Nitrogenase-molubdenum complex) | (Jack, De Zamaroczy, and Merrick 1999; Blanco et al. 1993) |
| YehUT | Nutrient Sensing | (Hirakawa, Nishino, Yamada, et al. 2003) |
| OmpR-Envz | Osmolarity stress, Acid pH Stress, biofilm formation, Type III fimbriae expression, | (T.-H. Lin et al. 2018; Bhagirath et al. 2019) |
| RcsCB | Regulated Phospho transfer, regulates Chlorpromazine-Induced Stress, biofilm formation | (Su et al. 2018) |
| PgtBA | Phosphoglycerate transport | (Niu, Jiang, and Hong 1995) |
| YpdAB | Pyruvate sensing | (Eguchi et al. 2004) |
| ArcBA | Regulates TCA cycle | (Ramos et al. 2016; Bhagirath et al. 2019) |
| TtrSR | Regulates Tetrathione reductase | (Chan, Kim, and Falkow 2005) |
| CitAB | Regulates Citrate fermentation | (Scheu et al. 2012; Y.-T. Chen et al. 2009) |
| TorSR | Regulates Trimethylamine-N-oxide sensing | (Toro-Roman, Wu, and Stock 2005) |
| ZraSR | Zinc Responsiveness | (Appia-Ayme et al. 2012) |

***Supplementary Table S4:*** *Detailed list of Two-component systems and their functions among Acinetobacter baumannii*

| **TCS (*A. baumannii*)** | **Function** | **References** |
|---|---|---|
| PmrBA | Colistin resistance, lipid modifications | (S. Sun et al. 2009; De Silva and Kumar 2019) |
| AdeSR | Controls expression of adeABC efflux pump involved in multidrug resistance | (Wen et al. 2017; M.-F. Lin et al. 2014) |
| BaeSR | modulate the expression of AdeIJK and MacAB-TolC efflux pumps, regulates osmotic stress under high osmotic stress | (De Silva and Kumar 2019) |
| KdpDE | Regulates Potassium sensing | (Samir et al. 2016) |
| CusSR | Regulates Copper sensing | (Alquethamy et al. 2019) |
| CzcSR | Heavy Metal sensing | (Chevalier et al. 2017) |
| IrlSR | Heavy metal ion resistance | (Lazar Adler et al. 2016) |
| CreCB | Regulates cells in fermenting glycolytic C source | (Nishino, Honda, and Yamaguchi 2005; Avison et al. 2001) |
| LytSR | Autolysin, regulates extracellular murein hydrolase activity, bacterial cell death and pyruvate utilization | (Radin et al. 2016) |
| PhoRB | Antibiotic Biosynthesis, phosphate regulation | (Bem et al. 2015) |
| RstBA | Responds to change in environmental condition, biofilm formation | (Zhou et al. 2003) |
| PilSR | Motility | (Richmond et al. 2016) |
| QseCB | Motility | (Bem et al. 2015) |
| GlnLG | Regulates Nitrogen regulation | (Ninfa and Magasanik 1986) |
| NtrBC | Regulates Nitrogen sensing | (Bhagirath et al. 2019) |
| OmpR-Envz | Osmolarity stress/Acid pH Stress | (Tipton and Rather 2017) |
| GacSR | Virulence, biofilm formation, pili formation | (Bhuiyan et al. 2016) |
| AlgZR | Virulence, twitching motility, alginate production | (Okkotsu, Little, and Schurr 2014) |

***Supplementary Table S5:*** *Detailed list of Two-component systems and their functions among Pseudomonas aeruginosa*

| **TCS (*P. aeruginosa)*** | **Function** | **References** |
|---|---|---|

| | | |
|---|---|---|
| CpxAR | Aminoglycoside resistance, Involves in mechanism of cell death | (Raivio 2014; Bem et al. 2015) |
| PhoQP | Polymyxin resistance, lipopolysaccharide modification | (Yamamoto et al. 2005; Francis, Stevenson, and Porter 2017) |
| CprSR | Antimicrobial peptide Resistance, Lipopolysaccharide modification | (Francis, Stevenson, and Porter 2017; Gooderham and Hancock 2009) |
| ErbR/EraR | Biofilm resistance antibiotic resistance | (Mern et al. 2010; Beaudoin et al. 2012) |
| PmrBA | Colistin resistance, antimicrobial peptide resistance | (Francis, Stevenson, and Porter 2017; Barbosa et al. 2017) |
| ParRS | Polymyxin resistance, quorum sensing, Lipopolysaccharide modification | (Muller, Plésiat, and Jeannot 2011; Barbosa et al. 2017) |
| KdpDE | Regulates potassium transport | (Freeman, Dorus, and Waterfield 2013) |
| PfeSR | Regulates iron acquisition | (Dean, Neshat, and Poole 1996) |
| KinB/AlgB | Regulates alginate biosynthesis for biofilm formation, quorum sensing, | (Chand et al. 2011; Chand, Clatworthy, and Hung 2012) |
| AruSR | Regulates arginine transaminase (ATA) pathway | (Bhagirath et al. 2019; C. Li, Yao, and Lu 2010) |
| MifSR | Regulates biofilm and metabolism, Alpha-ketoglutarate utilization | (Tatke et al. 2015; Petrova and Sauer 2010) |
| NarXL | Regulates biofilm and nitrogen sensing | (Francis, Stevenson, and Porter 2017) |
| BfiSR | Regulates biofilm formation and maintenance | (Petrova and Sauer 2010) |

| | | |
|---|---|---|
| BfmSR | Regulates biofilm formation and maintenance | (Petrova and Sauer 2010; Cerqueira et al. 2014) |
| BqsSR | Regulates biofilm decay | (Dong et al. 2008; Chevalier et al. 2017) |
| CarSR | Regulates biofilm decay, ferrous iron sensing | (Guragain et al. 2016) |
| DctBD | Regulates C4-dicarboxylate transporters | (Y.-T. Chen, Chang, Lu, et al. 2004; Sonawane, Singh, and Röhm 2006) |
| CbrAB | Controls Carbon & Nitrogen metabolism | (W. Li and Lu 2007) |
| CheAY | Regulates chemotaxis in response to magnesium | (Bhagirath et al. 2019) |
| TctED | Citrate metabolism under anaerobic condition | (Taylor, Zhang, and Mah 2019) |
| RoxSR | Confer cyanide tolerance | (Fernández-Piñar et al. 2008; Kawakami et al. 2010) |
| PvrSR | Regulates efflux pump and Biofilm formation | (Mikkelsen et al. 2013; Francis et al. 2018) |
| RocS2A2 | Regulates Fimbriae Adhesion | (Hurley et al. 2010) |
| FleSR | Regulates flagellar motility and adhesion to mucin | (Gellatly et al. 2018) |
| GtrSR | Controls Glucose transport and type III secretion | (Barbosa et al. 2017; Udaondo et al. 2018) |
| CzcSR | Regulates heavy metal resistance, quorum sensing, motility | (Chevalier et al. 2017) |
| PirRS | Regulates Iron acquisition and type IV pili | (Kilmury and Burrows 2018) |
| PprAB | Confer membrane permeability, assembling fimbriae | (Kilmury and Burrows 2018; Giraud et al. 2011) |
| QseCB | Motility, quorum sensing | (Tiwari et al. 2017; Bem et al. 2015) |
| NtrBC | Regulates nitrogen sensing | (W. Li and Lu 2007) |

| | | |
|---|---|---|
| OmpR-EnvZ | Regulates Osmolarity stress/Acid pH Stress | (Lau et al. 2013) |
| RegBA | Regulates Oxidative Respiration and Denitrification | (Willett and Crosson 2017) |
| RcsCB | Regulates biofilm formation, fimbriae, Phospho transfer | (Mikkelsen et al. 2009) |
| CreCB | Regulates cells in fermenting glycolytic C source, biofilm formation | (Zamorano et al. 2014) |
| HptBR | Regulates Swarming motility and Biofilm | (Hsu et al. 2008) |
| PhoRB | Regulates Swarming motility and quorum sensing | (Blus-Kadosh et al. 2013; Bains, Fernández, and Hancock 2012) |
| PilSR | Regulates Swarming/twitching motility and Biofilm | (Kilmury and Burrows 2018; Hobbs et al. 1993) |
| CopSR | Regulates tolerance to Copper and Zinc, imipenem resistance | (Caille, Rossier, and Perron 2007; Francis, Stevenson, and Porter 2017) |
| AlgZR | Differentially express virulence factors, sense environmental changes | (Okkotsu, Little, and Schurr 2014; Pritchett et al. 2015) |

**Supplementary Table S6:** Detailed list of Two-component systems and their functions among *Enterobacter cloacae*

| **TCS (*E. cloacae*)** | **Function** | **References** |
|---|---|---|
| CpxAR | Antibiotic resistance (fosfomycin), Carbon substrate uptake | (Roberts et al. 2011) |
| PhoQP | Polymyxin resistance, L-aminoarabinose synthesis regulation | (Kang et al. 2019) |
| PmrBA | Antibiotic resistance (Colistin), iron response, zinc response | (Kang et al. 2019) |
| BaeSR | Multidrug resistance, flagellar biosynthesis, maltose transport, chemotactic responses | (Roberts et al. 2011) |
| KdpDE | Regulates Potassium transport, intracellular survival of pathogenic bacteria | (Trastoy et al. 2018; Feeney and Sleator 2011) |
| CusSR | Regulates copper sensing and copper efflux | (Staehlin et al. 2016; Porcheron et al. 2013) |

| | | |
|---|---|---|
| PhoRB | Regulates Antibiotic Biosynthesis | (Knopp and Andersson 2015) |
| NarXL | Regulates Biofilm and Nitrogen sensing | (Huynh, Noriega, and Stewart 2010) |
| RstBA | Regulates change in environmental condition | (Hirakawa, Nishino, Hirata, et al. 2003) |
| TctED | Citrate metabolism under anaerobic condition | (Ren et al. 2010) |
| DcuSR | Regulates Fumarate respiration | (Abo-Amer et al. 2004; Zschiedrich, Keidel, and Szurmant 2016) |
| UhpBA | Regulates Hexose phosphate metabolism | (Yamamoto et al. 2005) |
| RcsCB | Regulates Indole Stress/ethanol/Nacl | (Yamamoto et al. 2005) |
| QseCB | Motility, quorum sensing | (Yamamoto et al. 2005) |
| NtrBC | Regulates Nitrogen sensing | (Nixon, Ronson, and Ausubel 1986) |
| OpmR-EnvZ | Regulates Osmolarity stress/Acid pH Stress | (Majewski et al. 2016) |
| YpdAB | Regulates Pyruvate sensing | (Behr, Brameyer, et al. 2017; Behr, Kristoficova, et al. 2017) |
| CopSR | Resistance to Copper | (Kremer and Hoffmann 2012; Staehlin et al. 2016) |
| BtsSR | Senses nutrient limitation | (Staehlin et al. 2016) |
| GlrKR | Stationary growth phase stimuli | (Keller et al. 1998) |
| BvgSA | Regulates virulence factor for colonization | (Lesne et al. 2016; Sobran and Cotter 2019) |

***Supplementary Table S7:*** *Detailed list of Two-component systems and their functions among Escherichia coli*

| **TCS (*E. coli*)** | **Function** | **References** |
|---|---|---|
| EvgSA | Multidrug resistance, Acid resistance | (Eguchi et al. 2004; |

| | | |
|---|---|---|
| | | Masuda and Church 2002) |
| BaeSR | Multidrug resistance, flagellar biosynthesis, maltose transport, chemotactic responses | (Nishino, Honda, and Yamaguchi 2005; Leblanc, Oates, and Raivio 2011) |
| BasSR/PmrBA | Antibiotic resistance (Colistin), iron response, zinc response | (Ogasawara et al. 2012; Folkesson et al. 2008) |
| CpxAR | Antibiotic resistance (fosfomycin), Carbon substrate uptake | (Kurabayashi et al. 2014) |
| CreCB | Others (Regulates cells in fermenting glycolytic C source) | (Amemura et al. 1986; Avison et al. 2001) |
| CusSR | Copper intake (copper related homeostasis) | (Ravikumar et al. 2012; Munson et al. 2000) |
| DpiBA/CitAB | Citrate metabolism under anaerobic condition/anaerobic condition | (Zhou et al. 2003) |
| GlrKR/QseEF | Envelop and LPS homeostasis, signal perception (Quorum sensing) | (Göpel et al. 2011) |
| KdpDE | K+ stimuli transport, Osmolarity stress, virulence, intracellular survival of pathogenic bacteria | (Jung et al. 2018; Freeman, Dorus, and Waterfield 2013) |
| NarXL | controls anaerobic respiratory gene expression in response to nitrate and nitrite | (Härtig et al. 1999; Landry et al. 2018) |
| NtrBC | Nitrogen regulation during N starve condition | (Brown et al. 2014) |
| OmpR-Envz | Osmolarity stress, Acid pH Stress | (Forst and Roberts 1994; Yuan et al. 2011) |
| PhoQP | Virulence, mediates the adaptation to Mg2+-limiting environments, and regulates numerous cellular activities | (Groisman 2001; Miyashiro and Goulian 2007) |
| PhoRB | Senses and respond extracellular inorganic phosphate | (Gao and Stock 2013; Hsieh and Wanner 2010) |
| QseCB | AI-3 quorum-sensing system (epinephrine/ nor-epinephrine) | (Furniss and Clements 2018; Tiwari et al. 2017) |

| | | |
|---|---|---|
| RcsCB | Others (Indole Stress/ethanol/Nacl), desiccation and osmotic shock | (Conter et al. 2002) |
| RstBA | Acid stress, curli fimbriae formation, and anaerobic respiration | (Yamamoto et al. 2005; Minagawa et al. 2003) |
| TorSR | Alternative anaerobic electron acceptors, alkaline stress defense mechanisms | (Schmidl et al. 2019; Ma et al. 2017) |
| UhpBA | Glucose -6-phosphate sensing | (Weston and Kadner 1988; Dahl, Wei, and Kadner 1997) |
| YedVW | Stress response to hydrogen peroxide | (Singh, Senadheera, and Cvitkovitch 2014; Urano et al. 2017) |
| YehUT/Bts SR | Nutrient Sensing regulatory network | (Kraxenberger et al. 2012) |
| ZraSR | Zinc sensing | (Petit-Härtlein et al. 2015; Rome et al. 2018) |
| DctBD | C4-dicarboxylate sensing | (Golby et al. 1999; Zientz, Bongaerts, and Unden 1998) |
| PgtBA | Phosphoglycerate transport | (Wanner 1992) |
| YpdAB/Pyr SR | Pyruvate sensing | (Miyake et al. 2019; Jung et al. 2018) |
| PcoSR | Copper-inducible expression of copper resistance | (Munson et al. 2000) |
| TtrSR | Tetrathione reductase | (Landry et al. 2018) |
| AtoSC | Acetoacetate sensing, motility, and chemotaxis | (Theodorou, Theodorou, and Kyriakidis 2012; Kyriakidis and Tiligada 2009) |
| DcuSR | Fumarate regulation | (Zientz, Bongaerts, and Unden 1998; Unden, Wörner, and Monzel 2016) |

### S2. Supplementary Figures

**A.**

| Pathogens | Genomes | QCQA1 | QCQA2 | QCQA3 | QCQA4 | QCQA5 |
|---|---|---|---|---|---|---|
| ***Enterococcus faecium*** | 1586 | 1417 | 1212 | 390 | 381 | 381 |
| ***Staphylococcus faecium*** | 8132 | 8132 | 208 | 1208 | 1166 | 1166 |
| ***Klebsiella pneumoniae*** | 7073 | 7073 | 2106 | 1189 | 1141 | 1141 |
| ***Acinetobacter baumannii*** | 3666 | 2648 | 1135 | 577 | 556 | 556 |
| ***Pseudomonas aeruginosa*** | 2308 | 2256 | 1771 | 969 | 929 | 929 |
| ***Enterobacter cloacae*** | 815 | 815 | 485 | 342 | 330 | 330 |
| ***Escherichia coli*** | 8554 | 8554 | 3711 | 1271 | 1226 | 1226 |

**B.** ***Enterococcus faecium***

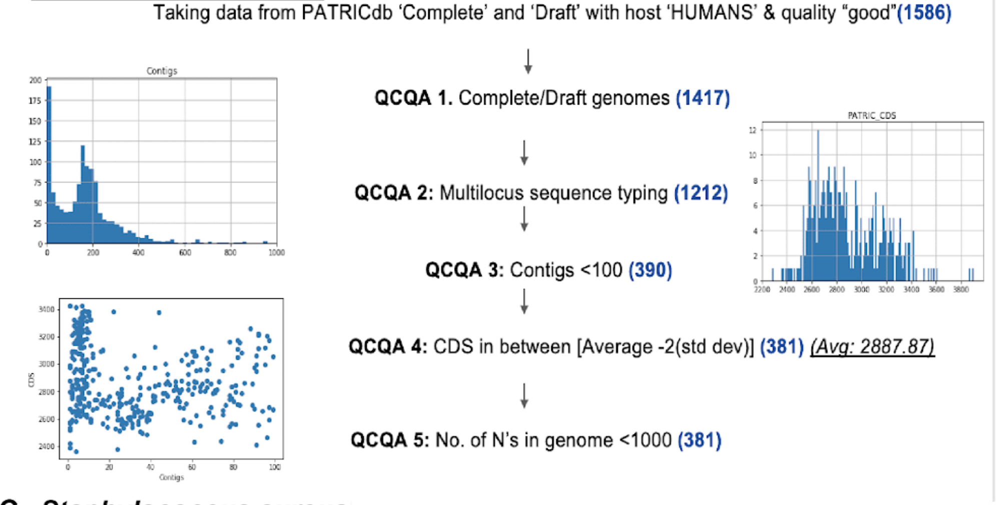


**C.** ***Staphylococcus aureus***

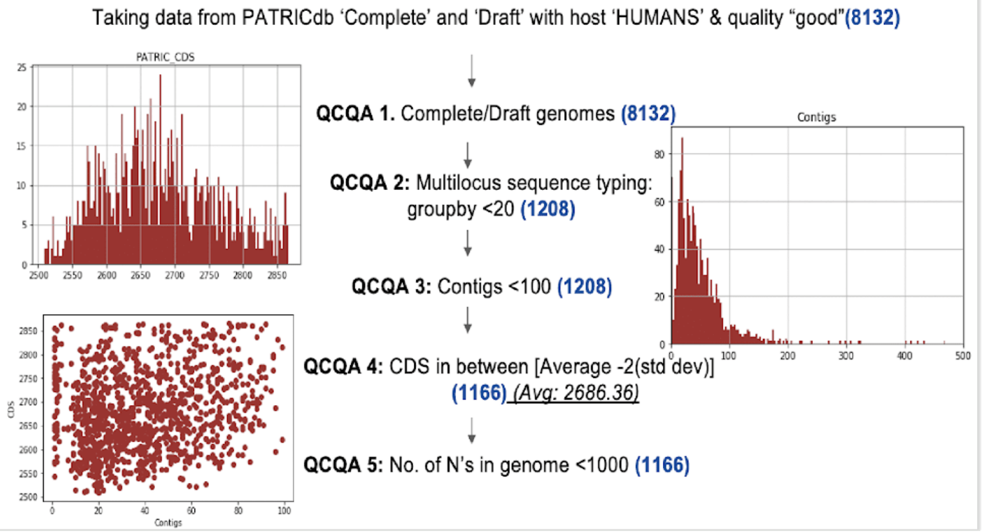


***Supplementary Figure S1:*** *Flow charts depicting the quality control steps and scatter plots for A) Table showing total number of genomes filtered at each step quality control steps, B) Enterococcus faecium, B) Staphylococcus aureus*

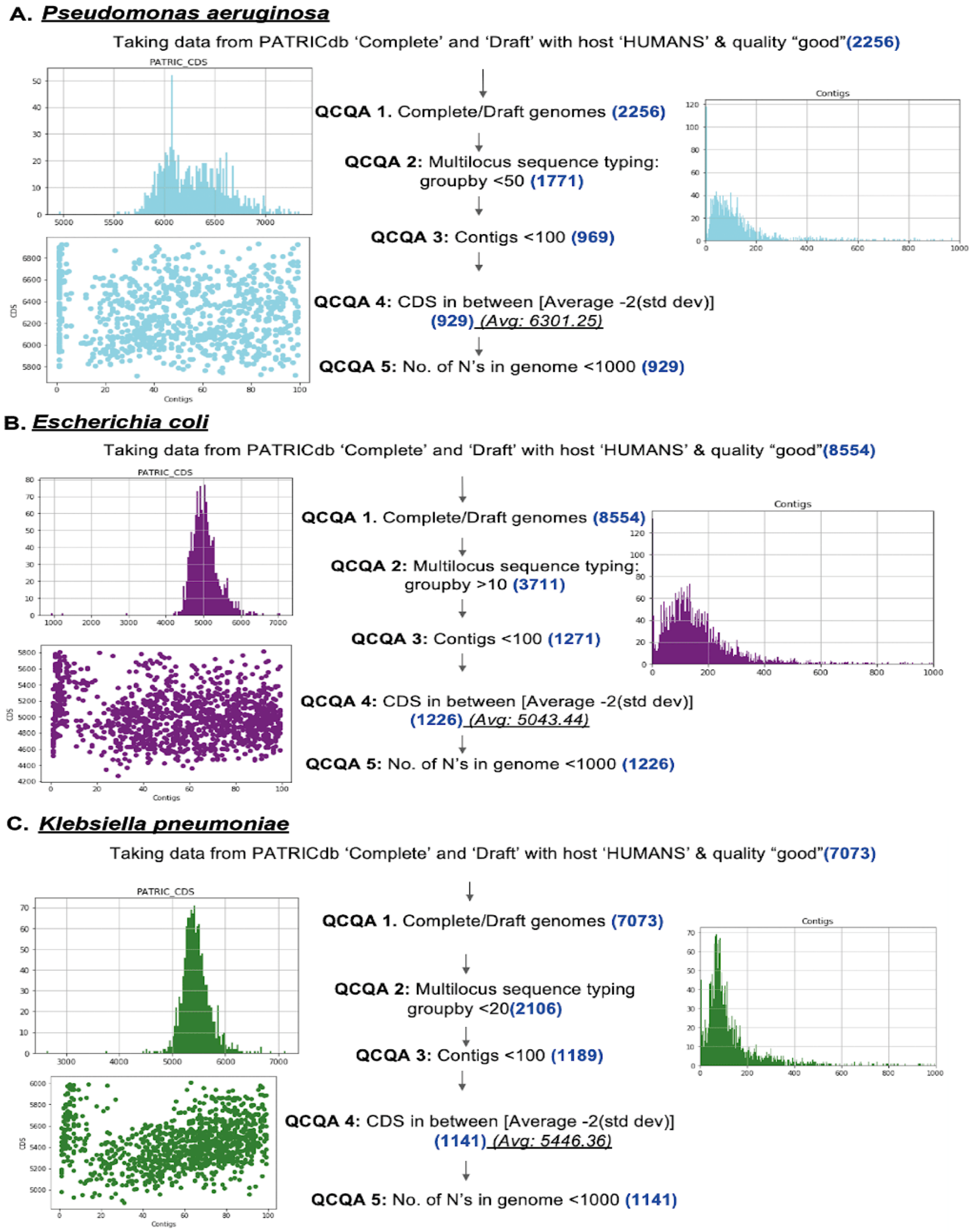


***Supplementary Figure S2:*** *Flow charts depicting the quality control steps and scatter plots for A) Pseudomonas aeruginosa, B) Escherichia coli, C) Klebsiella pneumoniae*

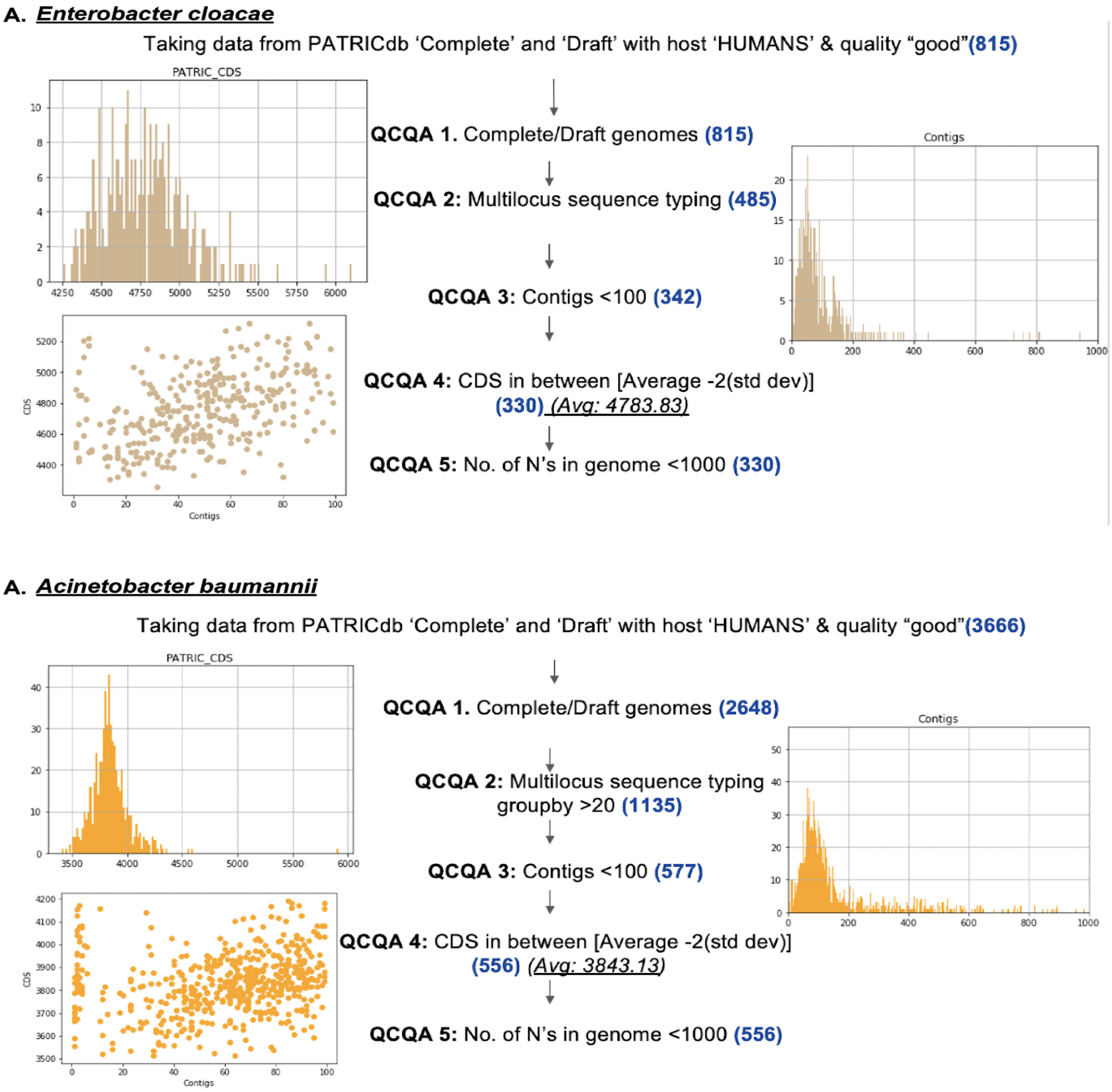


***Supplementary Figure S3:*** *Flow charts depicting the quality control steps and scatter plots for A) Enterobacter cloacae, B) Acinetobacter baumannii*

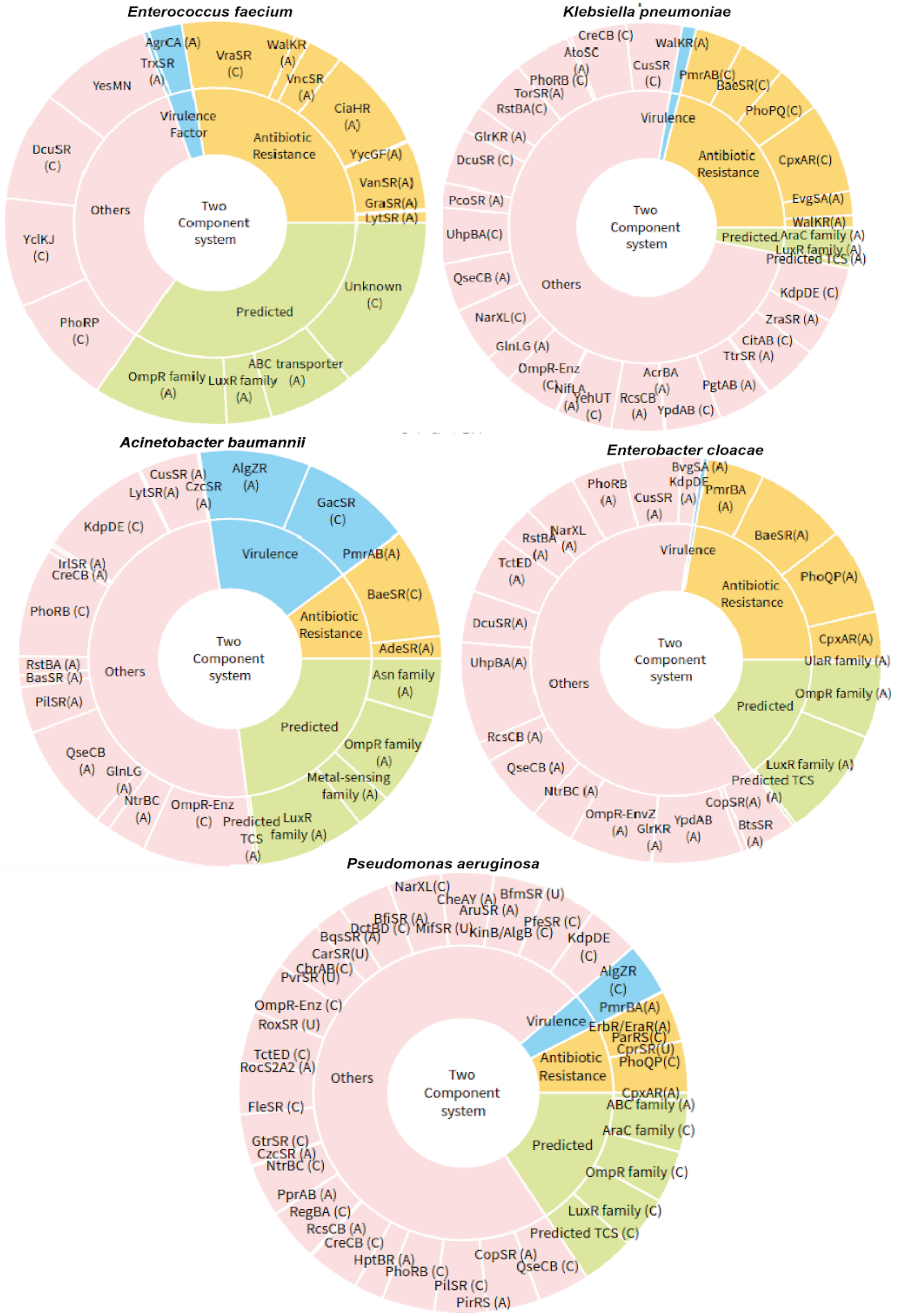

Enterococcus faecium
Two Component system
Virulence Factor
AgrCA (A)
TrxSR (A)
Antibiotic Resistance
VraSR (C)
WalKR (A)
VncSR (A)
CiaHR (A)
YycGF(A)
VanSR(A)
GraSR(A)
LytSR (A)
Predicted
Unknown (C)
ABC transporter (A)
LuxR family (A)
OmpR family (A)
Others
YesMN
DcuSR (C)
YclKJ (C)
PhoRP (C)
Klebsiella pneumoniae
Two Component system
Virulence
WalKR(A)
Antibiotic Resistance
PmrAB(C)
BaeSR(C)
PhoPQ(C)
CpxAR(C)
EvgSA(A)
WalKR(A)
Predicted
AraC family (A)
LuxR family (A)
Predicted TCS (A)
Others
CreCB (C)
AtoSC (A)
CusSR (C)
PhoRB (C)
TorSR(A)
RstBA(C)
GlrKR (A)
DcuSR (C)
PcoSR (A)
UhpBA(C)
QseCB (A)
NarXL(C)
GlnLG (A)
OmpR-Enz (C)
NifLA (A)
YehUT (C)
RcsCB (A)
AcrBA (A)
YpdAB (C)
PgtAB (A)
TtrSR (A)
CitAB (C)
ZraSR (A)
KdpDE (C)
Acinetobacter baumannii
Two Component system
Virulence
AlgZR (A)
GacSR (C)
Antibiotic Resistance
PmrAB(A)
BaeSR(C)
AdeSR(A)
Predicted
Asn family (A)
OmpR family (A)
Metal-sensing family (A)
LuxR family (A)
Predicted TCS (A)
Others
CusSR (A)
CzcSR (A)
LytSR(A)
KdpDE (C)
IrlSR (A)
CreCB (A)
PhoRB (C)
RstBA (A)
BasSR (A)
PilSR(A)
QseCB (A)
GlnLG (A)
NtrBC (A)
OmpR-Enz (C)
Enterobacter cloacae
Two Component system
Virulence
BvgSA (A)
KdpDE (A)
Antibiotic Resistance
PmrBA (A)
BaeSR(A)
PhoQP(A)
CpxAR(A)
Predicted
UlaR family (A)
OmpR family (A)
LuxR family (A)
Predicted TCS (A)
Others
PhoRB (A)
CusSR (A)
RstBA (A)
NarXL (A)
TctED (A)
DcuSR(A)
UhpBA(A)
RcsCB (A)
QseCB (A)
NtrBC (A)
OmpR-EnvZ (A)
GlrKR (A)
YpdAB (A)
CopSR(A)
BtsSR (A)
Pseudomonas aeruginosa
Two Component system
Virulence
AlgZR (C)
Antibiotic Resistance
PmrBA(A)
ErbR/EraR(A)
ParRS(C)
CprSR(U)
PhoQP(C)
CpxAR(A)
Predicted
ABC family (A)
AraC family (C)
OmpR family (C)
LuxR family (C)
Predicted TCS (C)
Others
NarXL(C)
CheAY (A)
BfmSR (U)
AruSR (A)
BfiSR (A)
DctBD (C)
MifSR (U)
KinB/AlgB (C)
PfeSR (C)
KdpDE (C)
BqsSR (A)
CarSR(U)
ChrAB(C)
PvrSR (U)
OmpR-Enz (C)
RoxSR (U)
TctED (C)
RocS2A2 (A)
FleSR (C)
GtrSR (C)
CzcSR (A)
NtrBC (C)
PprAB (A)
RegBA (C)
RcsCB (A)
CreCB (C)
HptBR (A)
PhoRB (C)
PilSR (C)
PirRS (A)
CopSR (A)
QseCB (C)

***Supplementary Figure S4:*** *Pan genome analysis of the two-component systems in form of multilevel pie chart depicting the distribution of TCS in all the four categories in Enterococcus faecium, Klebsiella pneumoniae, Acinetobacter baumannii, and Enterobacter cloacae, and Pseudomonas aeruginosa*

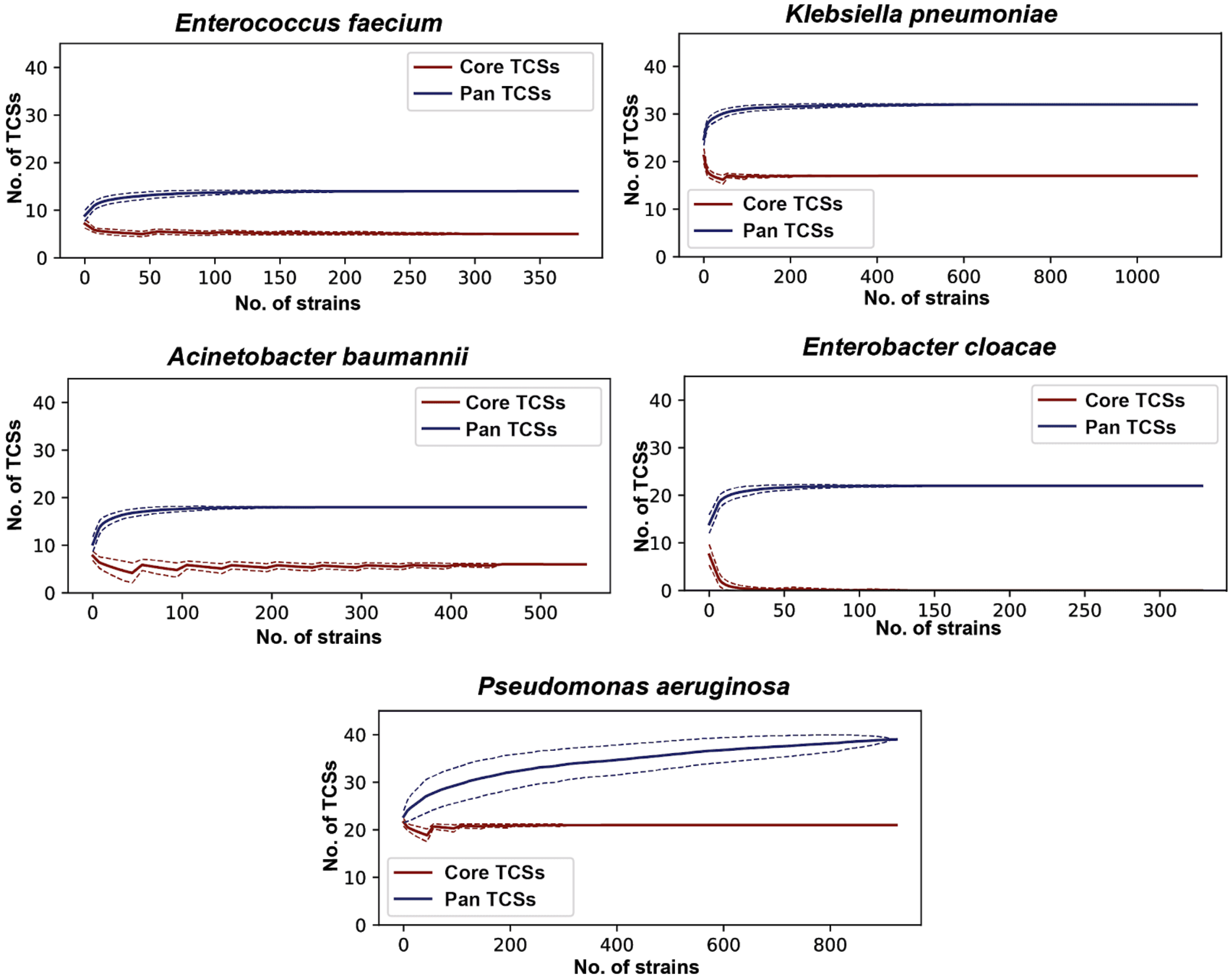


***Supplementary Figure S5:*** *Pan genome analysis of the two-component systems in form of the Pangenome curve of Enterococcus faecium, Klebsiella pneumoniae, Acinetobacter baumannii, and Enterobacter cloacae, and Pseudomonas aeruginosa. The curve shows the status of core and pan-genome for TCS. All the four pathogens show "closed" pangenome for TCS except P. aeruginosa*

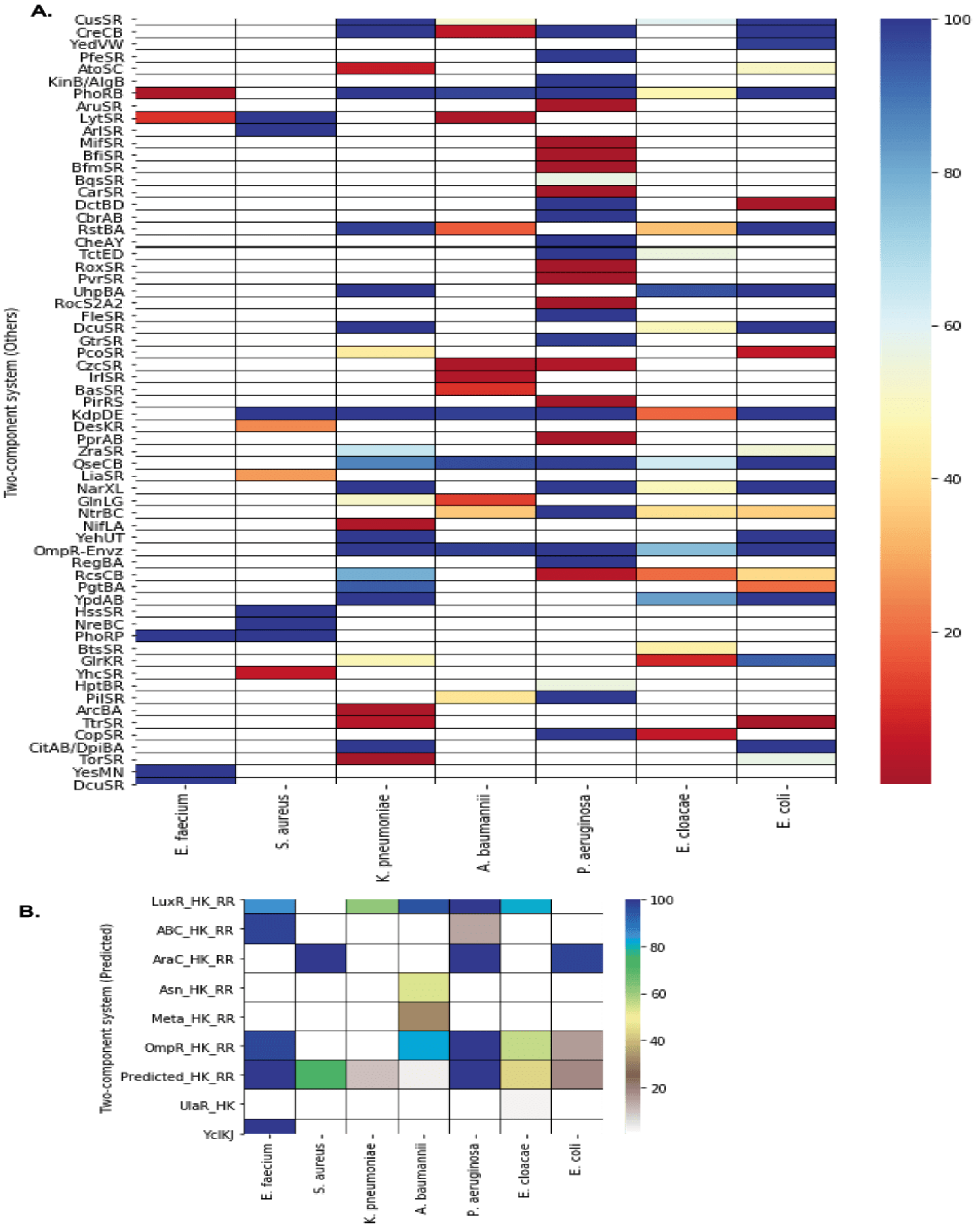


***Supplementary Figure S6:*** *Pan genome analysis of the two-component systems: A) Heatmaps depicting the TCS involved in others (general) category, B) Heatmaps depicting the predicted family of TCS*

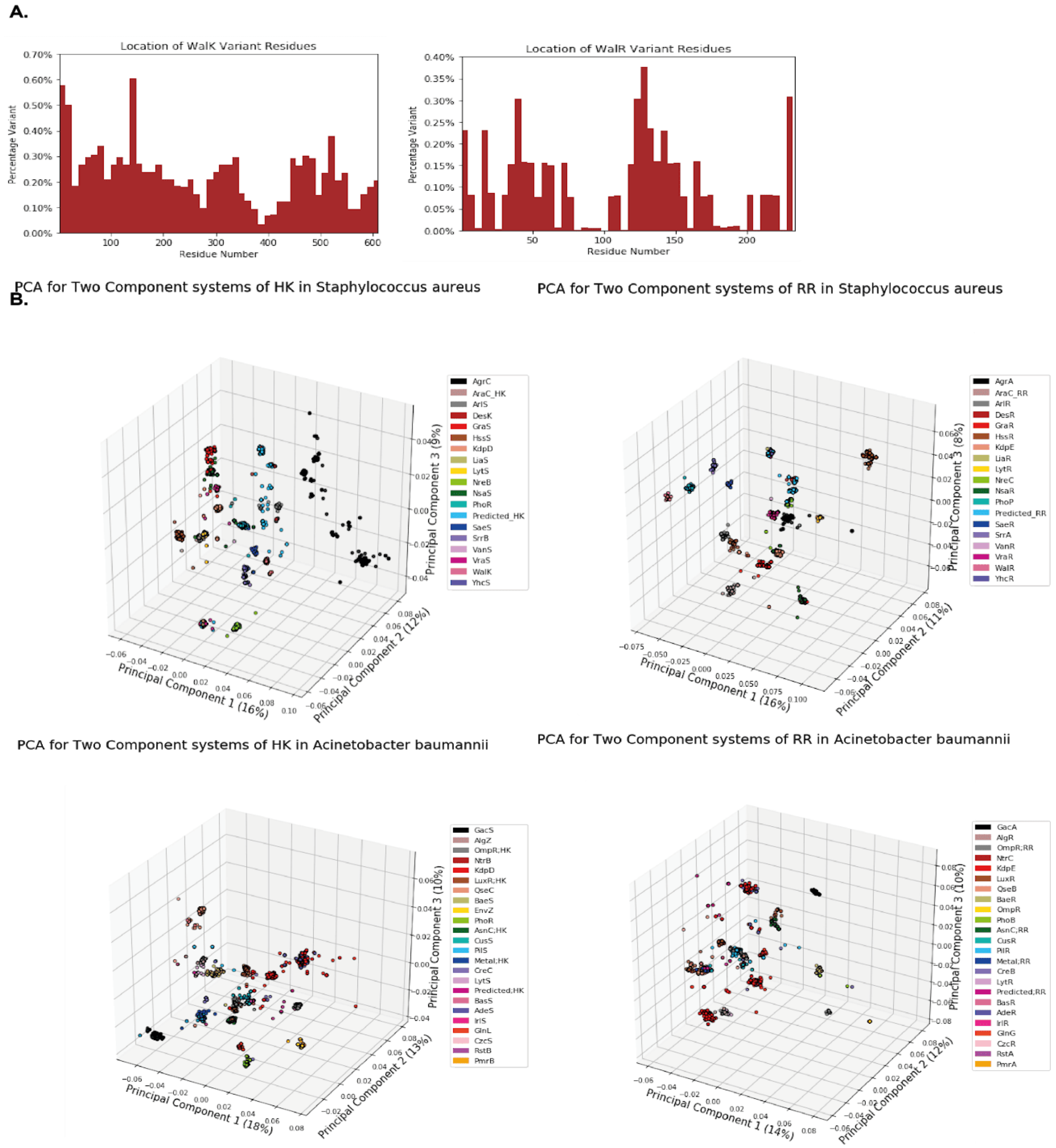


***Supplementary Figure S7:*** *Pan genome analysis of the two-component systems: A) Sequence variants bar graphs of WalK and WalR TCS. The graph is plotted between percentage variation v/s Number of residues, B) Principal component Analysis (PCA) curves showing histidine kinase and response regulators of Staphylococcus aureus and Acinetobacter baumannii. The PCA curves plotted by using peptide features i.e. amino acid composition, dipeptide composition, and tripeptide composition*

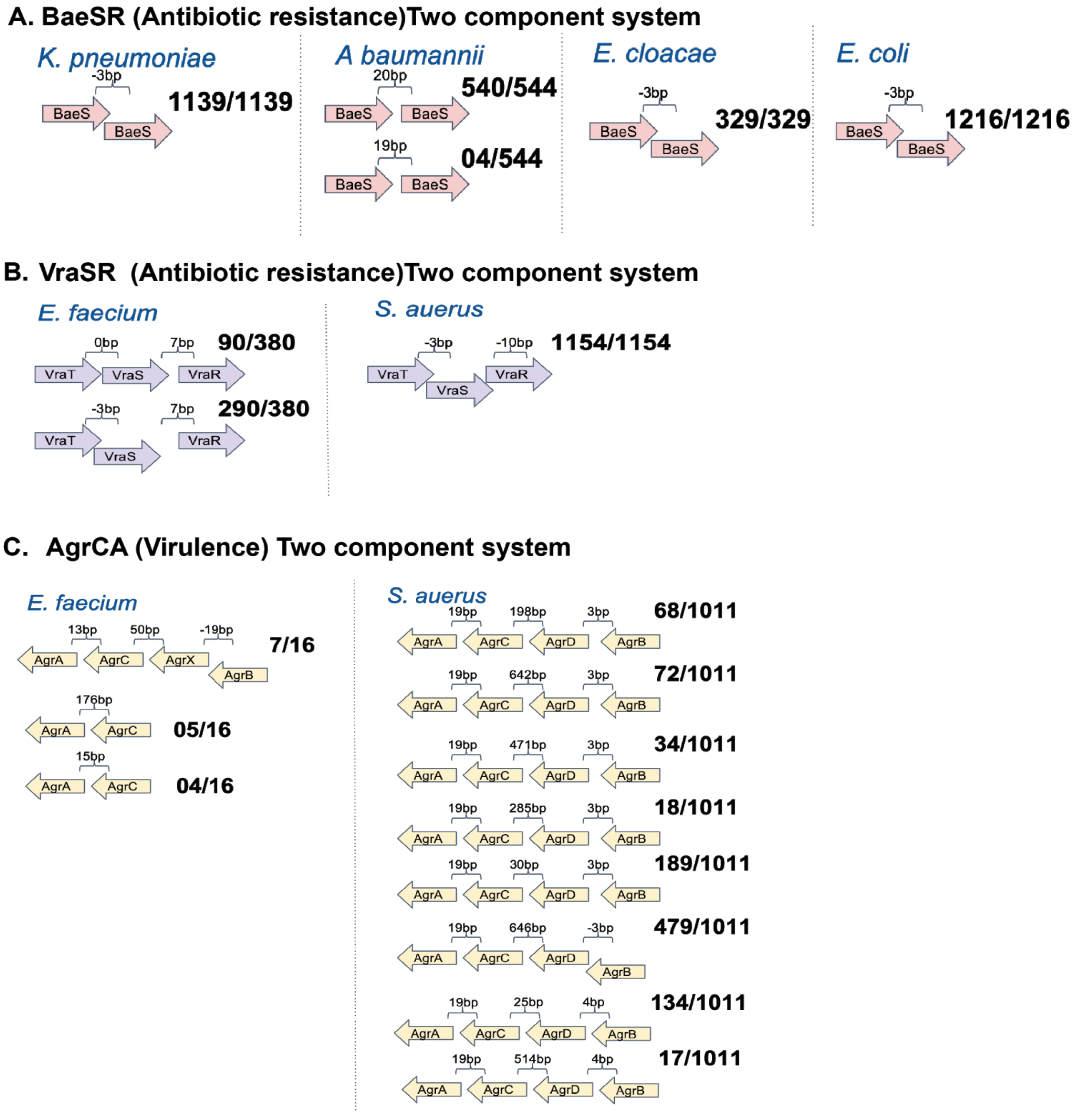


***Supplementary Figure S8:*** *Pan genome analysis of the two-component systems: A) Genomic architecture of BaeSR antibiotic resistance two-component system among Gram-negative ESKAPEE pathogens i.e. K. pneumoniae, A. baumannii, E. cloacae, and E. coli. B) Genomic architecture of VraSR antibiotic resistance two-component system. The VraSR system found in Gram-positive ESKAPEE pathogens i.e. E. faecium, and S. aureus. C) Genomic architecture of AgrCA virulence two-component system in E. faecium, and S. aureus*

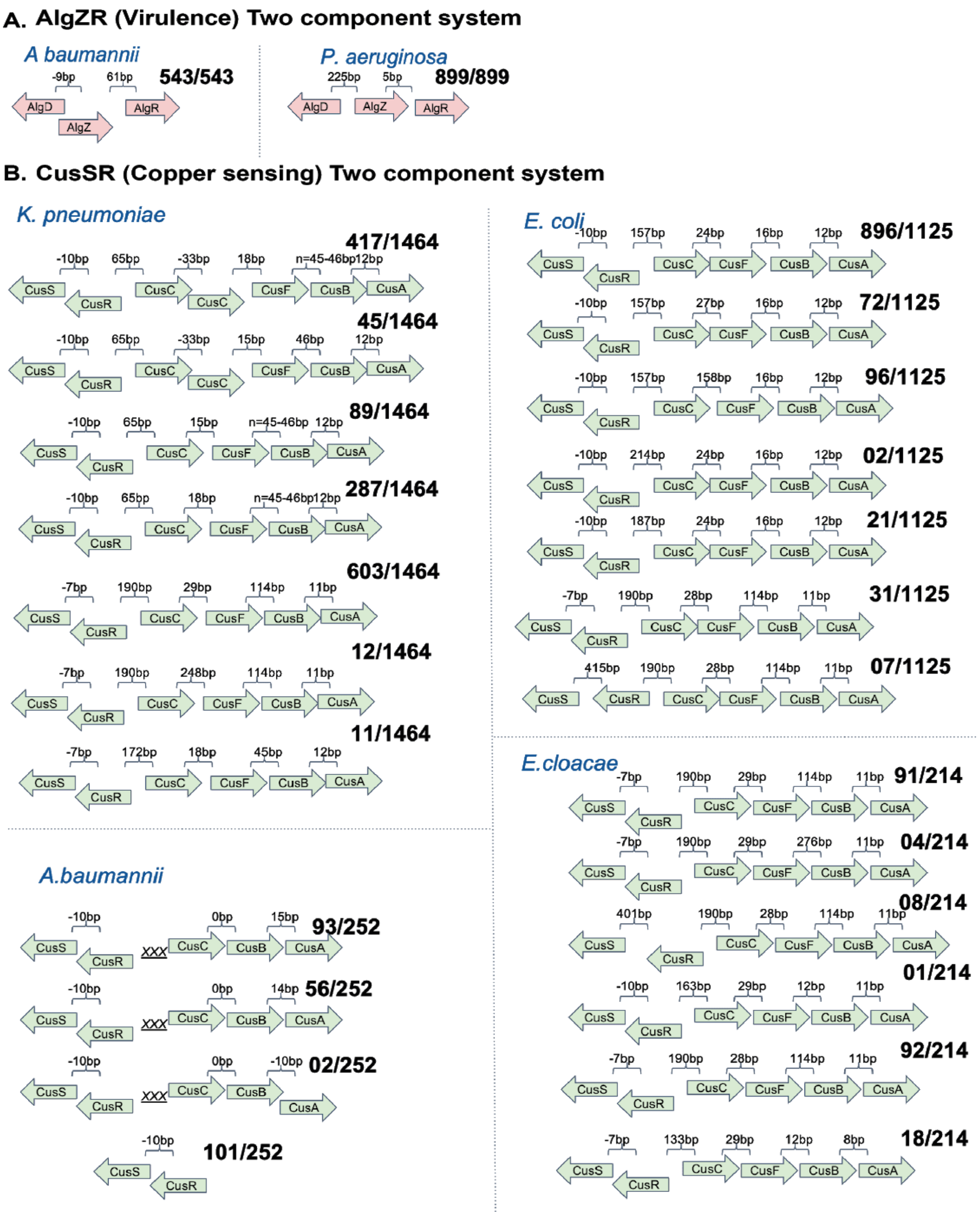


***Supplementary Figure S9:*** *Pan genome analysis of the two-component systems: A) Genomic architecture of AlgZR virulence two-component system among Gram-negative ESKAPEE pathogens i.e. A. baumannii, and P .aeruginosa. B) Genomic architecture of CusSR Copper sensing two-component system. The CusSR system found in K. pneumoniae, A. baumannii, E. cloacae, and E. coli*

Armstrong, Brent D., Christine A. Herfst, Nicholas C. Tonial, Adrienne T. Wakabayashi, Joseph J. Zeppa, and John K. McCormick. 2016. “Identification of a Two-Component Class IIb Bacteriocin in Streptococcus Pyogenes by Recombinase-Based in Vivo Expression Technology.” *Scientific Reports* 6 (November): 36233.

Arthur, M., C. Molinas, and P. Courvalin. 1992. “The VanS-VanR Two-Component Regulatory System Controls Synthesis of Depsipeptide Peptidoglycan Precursors in Enterococcus Faecium BM4147.” *Journal of Bacteriology* 174 (8): 2582–91.

Avison, M. B., R. E. Horton, T. R. Walsh, and P. M. Bennett. 2001. “Escherichia Coli CreBC Is a Global Regulator of Gene Expression That Responds to Growth in Minimal Media.” *The Journal of Biological Chemistry* 276 (29): 26955–61.

Bains, Manjeet, Lucía Fernández, and Robert E. W. Hancock. 2012. “Phosphate Starvation Promotes Swarming Motility and Cytotoxicity of Pseudomonas Aeruginosa.” *Applied and Environmental Microbiology* 78 (18): 6762–68.

Barbosa, Camilo, Vincent Trebosc, Christian Kemmer, Philip Rosenstiel, Robert Beardmore, Hinrich Schulenburg, and Gunther Jansen. 2017. “Alternative Evolutionary Paths to Bacterial Antibiotic Resistance Cause Distinct Collateral Effects.” *Molecular Biology and Evolution* 34 (9): 2229–44.

Beaudoin, Trevor, Li Zhang, Aaron J. Hinz, Christopher J. Parr, and Thien-Fah Mah. 2012. “The Biofilm-Specific Antibiotic Resistance Gene ndvB Is Important for Expression of Ethanol Oxidation Genes in Pseudomonas Aeruginosa Biofilms.” *Journal of Bacteriology* 194 (12): 3128–36.

Behr, Stefan, Sophie Brameyer, Michael Witting, Philipp Schmitt-Kopplin, and Kirsten Jung. 2017. “Comparative Analysis of LytS/LytTR-Type Histidine Kinase/response Regulator Systems in γ-Proteobacteria.” *PloS One* 12 (8): e0182993.

Behr, Stefan, Ivica Kristoficova, Michael Witting, Erin J. Breland, Allison R. Eberly, Corinna Sachs, Philippe Schmitt-Kopplin, Maria Hadjifrangiskou, and Kirsten Jung. 2017. “Identification of a High-Affinity Pyruvate Receptor in Escherichia Coli.” *Scientific Reports* 7 (1): 1388.

Belcheva, Antoaneta, and Dasantila Golemi-Kotra. 2008. “A Close-up View of the VraSR Two-Component System. A Mediator of Staphylococcus Aureus Response to Cell Wall Damage.” *The Journal of Biological Chemistry* 283 (18): 12354–64.

Bem, Agnieszka E., Nadya Velikova, M. Teresa Pellicer, Peter van Baarlen, Alberto Marina, and Jerry M. Wells. 2015. “Bacterial Histidine Kinases as Novel Antibacterial Drug Targets.” *ACS Chemical Biology* 10 (1): 213–24.

Bhagirath, Anjali Y., Yanqi Li, Rakesh Patidar, Katherine Yerex, Xiaoxue Ma, Ayush Kumar, and Kangmin Duan. 2019. “Two Component Regulatory Systems and Antibiotic Resistance in Gram-Negative Pathogens.” *International Journal of Molecular Sciences* 20 (7). https://doi.org/10.3390/ijms20071781.

Bhuiyan, Md Saruar, Felix Ellett, Gerald L. Murray, Xenia Kostoulias, Gustavo M. Cerqueira, Keith E. Schulze, Mohd Hafidz Mahamad Maifiah, et al. 2016. “Acinetobacter Baumannii Phenylacetic Acid Metabolism Influences Infection Outcome through a Direct Effect on Neutrophil Chemotaxis.” *Proceedings of the National Academy of Sciences of the United States of America* 113 (34): 9599–9604.

Blanco, G., M. Drummond, P. Woodley, and C. Kennedy. 1993. “Sequence and Molecular Analysis of the nifL Gene of Azotobacter Vinelandii.” *Molecular Microbiology* 9 (4): 869–79.

Blus-Kadosh, Inna, Anat Zilka, Gal Yerushalmi, and Ehud Banin. 2013. “The Effect of pstS and phoB on Quorum Sensing and Swarming Motility in Pseudomonas Aeruginosa.” *PloS One* 8 (9): e74444.

Botella, Eric, Susanne Krogh Devine, Sebastian Hubner, Letal I. Salzberg, Robert T. Gale, Eric D. Brown, Hannes Link, et al. 2014. “PhoR Autokinase Activity Is Controlled by an Intermediate in Wall Teichoic Acid Metabolism That Is Sensed by the Intracellular PAS Domain during the PhoPR-Mediated Phosphate Limitation Response of Bacillus Subtilis.” *Molecular Microbiology* 94 (6): 1242–59.

Bronner, Stéphane, Henri Monteil, and Gilles Prévost. 2004. “Regulation of Virulence Determinants in Staphylococcus Aureus: Complexity and Applications.” *FEMS Microbiology Reviews* 28 (2): 183–200.

Brown, Daniel R., Geraint Barton, Zhensheng Pan, Martin Buck, and Sivaramesh Wigneshweraraj. 2014. “Nitrogen Stress Response and Stringent Response Are Coupled in Escherichia Coli.” *Nature Communications* 5 (June): 4115.

Burnside, Kellie, Annalisa Lembo, Melissa de Los Reyes, Anton Iliuk, Nguyen-Thao Binhtran, James E. Connelly, Wan-Jung Lin, et al. 2010. “Regulation of Hemolysin Expression and Virulence of Staphylococcus Aureus by a Serine/threonine Kinase and Phosphatase.” *PloS One* 5 (6): e11071.

Caille, Olivier, Claude Rossier, and Karl Perron. 2007. “A Copper-Activated

Two-Component System Interacts with Zinc and Imipenem Resistance in Pseudomonas Aeruginosa.” *Journal of Bacteriology* 189 (13): 4561–68.
Cerqueira, Gustavo M., Xenia Kostoulias, Chen Khoo, Ibukun Aibinu, Yue Qu, Ana Traven, and Anton Y. Peleg. 2014. “A Global Virulence Regulator in Acinetobacter Baumannii and Its Control of the Phenylacetic Acid Catabolic Pathway.” *The Journal of Infectious Diseases* 210 (1): 46–55.
Chand, Nikhilesh S., Anne E. Clatworthy, and Deborah T. Hung. 2012. “The Two-Component Sensor KinB Acts as a Phosphatase to Regulate Pseudomonas Aeruginosa Virulence.” *Journal of Bacteriology* 194 (23): 6537–47.
Chand, Nikhilesh S., Jenny See-Wai Lee, Anne E. Clatworthy, Aaron J. Golas, Roger S. Smith, and Deborah T. Hung. 2011. “The Sensor Kinase KinB Regulates Virulence in Acute Pseudomonas Aeruginosa Infection.” *Journal of Bacteriology* 193 (12): 2989–99.
Chan, Kaman, Charles C. Kim, and Stanley Falkow. 2005. “Microarray-Based Detection of Salmonella Enterica Serovar Typhimurium Transposon Mutants That Cannot Survive in Macrophages and Mice.” *Infection and Immunity* 73 (9): 5438–49.
Chen, Sheng-Chia, Chi-Fon Chang, Pei-Ju Fan, Ya-Hsin Cheng, Tsunai Yu, and Tai-Huang Huang. 2013. “(1)H, (13)C and (15)N Resonance Assignments of the C-Terminal DNA-Binding Domain of RstA Protein from Klebsiella Pneumoniae.” *Biomolecular NMR Assignments* 7 (1): 85–88.
Chen, Ying-Tsong, Hwan-You Chang, Yi-Chyi Lai, Chao-Chi Pan, Shih-Feng Tsai, and Hwei-Ling Peng. 2004. “Sequencing and Analysis of the Large Virulence Plasmid pLVPK of Klebsiella Pneumoniae CG43.” *Gene* 337 (August): 189–98.
Chen, Ying-Tsong, Hwan You Chang, Chin Lung Lu, and Hwei-Ling Peng. 2004. “Evolutionary Analysis of the Two-Component Systems in Pseudomonas Aeruginosa PAO1.” *Journal of Molecular Evolution* 59 (6): 725–37.
Chen, Ying-Tsong, Tsai-Lien Liao, Keh-Ming Wu, Tsai-Ling Lauderdale, Jing-Jou Yan, I-Wen Huang, Min-Chi Lu, et al. 2009. “Genomic Diversity of Citrate Fermentation in Klebsiella Pneumoniae.” *BMC Microbiology* 9 (August): 168.
Chevalier, Sylvie, Emeline Bouffartigues, Josselin Bodilis, Olivier Maillot, Olivier Lesouhaitier, Marc G. J. Feuilloley, Nicole Orange, Alain Dufour, and Pierre Cornelis. 2017. “Structure, Function and Regulation of Pseudomonas Aeruginosa Porins.” *FEMS Microbiology Reviews* 41 (5): 698–722.
Choudhary, Kumari S., Nathan Mih, Jonathan Monk, Erol Kavvas, James T. Yurkovich, George Sakoulas, and Bernhard O. Palsson. 2018. “The Two-Component System AgrAC Displays Four Distinct Genomic Arrangements That Delineate Genomic Virulence Factor Signatures.” *Frontiers in Microbiology* 9 (May): 1082.
Christmas, Benjamin A. F., Matthew D. Rolfe, Matthew Rose, and Jeffrey Green. 2019. “Staphylococcus Aureus Adaptation to Aerobic Low-Redox-Potential Environments: Implications for an Intracellular Lifestyle.” *Microbiology* 165 (7): 779–91.
Conter, Annie, Rachel Sturny, Claude Gutierrez, and Kaymeuang Cam. 2002. “The RcsCB His-Asp Phosphorelay System Is Essential to Overcome Chlorpromazine-Induced Stress in Escherichia Coli.” *Journal of Bacteriology* 184 (10): 2850–53.
Dahl, J. L., B. Y. Wei, and R. J. Kadner. 1997. “Protein Phosphorylation Affects Binding

of the Escherichia Coli Transcription Activator UhpA to the uhpT Promoter.” *The Journal of Biological Chemistry* 272 (3): 1910–19.

Dean, C. R., S. Neshat, and K. Poole. 1996. “PfeR, an Enterobactin-Responsive Activator of Ferric Enterobactin Receptor Gene Expression in Pseudomonas Aeruginosa.” *Journal of Bacteriology* 178 (18): 5361–69.

Delauné, Aurélia, Sarah Dubrac, Charlène Blanchet, Olivier Poupel, Ulrike Mäder, Aurélia Hiron, Aurélie Leduc, et al. 2012. “The WalKR System Controls Major Staphylococcal Virulence Genes and Is Involved in Triggering the Host Inflammatory Response.” *Infection and Immunity* 80 (10): 3438–53.

De Silva, P. Malaka, and Ayush Kumar. 2019. “Signal Transduction Proteins in Acinetobacter Baumannii: Role in Antibiotic Resistance, Virulence, and Potential as Drug Targets.” *Frontiers in Microbiology* 10 (January): 49.

Diaz, Lorena, Truc T. Tran, Jose M. Munita, William R. Miller, Sandra Rincon, Lina P. Carvajal, Aye Wollam, et al. 2014. “Whole-Genome Analyses of Enterococcus Faecium Isolates with Diverse Daptomycin MICs.” *Antimicrobial Agents and Chemotherapy* 58 (8): 4527–34.

Dong, Yi-Hu, Xi-Fen Zhang, Shu-Wen An, Jin-Ling Xu, and Lian-Hui Zhang. 2008. “A Novel Two-Component System BqsS-BqsR Modulates Quorum Sensing-Dependent Biofilm Decay in Pseudomonas Aeruginosa.” *Communicative & Integrative Biology* 1 (1): 88–96.

Dubrac, Sarah, Paola Bisicchia, Kevin M. Devine, and Tarek Msadek. 2008. “A Matter of Life and Death: Cell Wall Homeostasis and the WalKR (YycGF) Essential Signal Transduction Pathway.” *Molecular Microbiology* 70 (6): 1307–22.

Dubrac, Sarah, Ivo Gomperts Boneca, Olivier Poupel, and Tarek Msadek. 2007. “New Insights into the WalK/WalR (YycG/YycF) Essential Signal Transduction Pathway Reveal a Major Role in Controlling Cell Wall Metabolism and Biofilm Formation in Staphylococcus Aureus.” *Journal of Bacteriology* 189 (22): 8257–69.

Eguchi, Yoko, Tadashi Okada, Shu Minagawa, Taku Oshima, Hirotada Mori, Kaneyoshi Yamamoto, Akira Ishihama, and Ryutaro Utsumi. 2004. “Signal Transduction Cascade between EvgA/EvgS and PhoP/PhoQ Two-Component Systems of Escherichia Coli.” *Journal of Bacteriology* 186 (10): 3006–14.

Fakhruzzaman, Md, Yoichi Inukai, Yohei Yanagida, Hirokazu Kino, Masayuki Igarashi, Yoko Eguchi, and Ryutaro Utsumi. 2015. “Study on in Vivo Effects of Bacterial Histidine Kinase Inhibitor, Waldiomycin, in Bacillus Subtilis and Staphylococcus Aureus.” *The Journal of General and Applied Microbiology* 61 (5): 177–84.

Feeney, Audrey, and Roy D. Sleator. 2011. “An in Silico Analysis of Osmotolerance in the Emerging Gastrointestinal Pathogen Cronobacter Sakazakii.” *Bioengineered Bugs* 2 (5): 260–70.

Fernández-Piñar, Regina, Juan Luis Ramos, José Juan Rodríguez-Herva, and Manuel Espinosa-Urgel. 2008. “A Two-Component Regulatory System Integrates Redox State and Population Density Sensing in Pseudomonas Putida.” *Journal of Bacteriology* 190 (23): 7666–74.

Folkesson, Anders, Janus A. J. Haagensen, Claudia Zampaloni, Claus Sternberg, and Søren Molin. 2008. “Biofilm Induced Tolerance towards Antimicrobial Peptides.” *PloS One* 3 (4): e1891.

Forst, S. A., and D. L. Roberts. 1994. “Signal Transduction by the EnvZ-OmpR Phosphotransfer System in Bacteria.” *Research in Microbiology* 145 (5-6): 363–73.

Francis, Vanessa I., Emma C. Stevenson, and Steven L. Porter. 2017. “Two-Component Systems Required for Virulence in Pseudomonas Aeruginosa.” *FEMS Microbiology Letters* 364 (11). https://doi.org/10.1093/femsle/fnx104.

Francis, Vanessa I., Elaine M. Waters, Sutharsan E. Finton-James, Andrea Gori, Aras Kadioglu, Alan R. Brown, and Steven L. Porter. 2018. “Multiple Communication Mechanisms between Sensor Kinases Are Crucial for Virulence in Pseudomonas Aeruginosa.” *Nature Communications* 9 (1): 2219.

Freeman, Zoë N., Steve Dorus, and Nicholas R. Waterfield. 2013. “The KdpD/KdpE Two-Component System: Integrating $K^+$ Homeostasis and Virulence.” *PLoS Pathogens* 9 (3): e1003201.

Furniss, R. Christopher D., and Abigail Clements. 2018. “Regulation of the Locus of Enterocyte Effacement in Attaching and Effacing Pathogens.” *Journal of Bacteriology* 200 (2). https://doi.org/10.1128/JB.00336-17.

Gao, Rong, and Ann M. Stock. 2013. “Probing Kinase and Phosphatase Activities of Two-Component Systems in Vivo with Concentration-Dependent Phosphorylation Profiling.” *Proceedings of the National Academy of Sciences of the United States of America* 110 (2): 672–77.

Gellatly, Shaan L., Manjeet Bains, Elena B. M. Breidenstein, Janine Strehmel, Fany Reffuveille, Patrick K. Taylor, Amy T. Y. Yeung, Joerg Overhage, and Robert E. W. Hancock. 2018. “Novel Roles for Two-Component Regulatory Systems in Cytotoxicity and Virulence-Related Properties in.” *AIMS Microbiology* 4 (1): 173–91.

Giraud, Caroline, Christophe S. Bernard, Virginie Calderon, Liang Yang, Alain Filloux, Søren Molin, Gwennaele Fichant, Christophe Bordi, and Sophie de Bentzmann. 2011. “The PprA-PprB Two-Component System Activates CupE, the First Non-Archetypal Pseudomonas Aeruginosa Chaperone-Usher Pathway System Assembling Fimbriae.” *Environmental Microbiology* 13 (3): 666–83.

Giraudo, A. T., A. Calzolari, A. A. Cataldi, C. Bogni, and R. Nagel. 1999. “The Sae Locus of Staphylococcus Aureus Encodes a Two-Component Regulatory System.” *FEMS Microbiology Letters* 177 (1): 15–22.

Golby, P., S. Davies, D. J. Kelly, J. R. Guest, and S. C. Andrews. 1999. “Identification and Characterization of a Two-Component Sensor-Kinase and Response-Regulator System (DcuS-DcuR) Controlling Gene Expression in Response to C4-Dicarboxylates in Escherichia Coli.” *Journal of Bacteriology* 181 (4): 1238–48.

Gooderham, W. James, and Robert E. W. Hancock. 2009. “Regulation of Virulence and Antibiotic Resistance by Two-Component Regulatory Systems in Pseudomonas Aeruginosa.” *FEMS Microbiology Reviews* 33 (2): 279–94.

Göpel, Yvonne, Denise Lüttmann, Ann Kathrin Heroven, Birte Reichenbach, Petra Dersch, and Boris Görke. 2011. “Common and Divergent Features in Transcriptional Control of the Homologous Small RNAs GlmY and GlmZ in Enterobacteriaceae.” *Nucleic Acids Research* 39 (4): 1294–1309.

Groisman, E. A. 2001. “The Pleiotropic Two-Component Regulatory System PhoP-PhoQ.” *Journal of Bacteriology* 183 (6): 1835–42.

Guragain, Manita, Michelle M. King, Kerry S. Williamson, Ailyn C. Pérez-Osorio, Tatsuya Akiyama, Sharmily Khanam, Marianna A. Patrauchan, and Michael J. Franklin. 2016. “The Pseudomonas Aeruginosa PAO1 Two-Component Regulator CarSR Regulates Calcium Homeostasis and Calcium-Induced Virulence Factor Production through Its Regulatory Targets CarO and CarP.” *Journal of Bacteriology* 198 (6): 951–63.

Hancock, Lynn E., and Marta Perego. 2004. “The Enterococcus Faecalis Fsr Two-Component System Controls Biofilm Development through Production of Gelatinase.” *Journal of Bacteriology* 186 (17): 5629–39.

Hancock, Lynn, and Marta Perego. 2002. “Two-Component Signal Transduction in Enterococcus Faecalis.” *Journal of Bacteriology* 184 (21): 5819–25.

Hanczvikkel, Adrienn, Miklós Füzi, Erika Ungvári, and Ákos Tóth. 2018. “Transmissible Silver Resistance Readily Evolves in High-Risk Clone Isolates of Klebsiella Pneumoniae.” *Acta Microbiologica et Immunologica Hungarica* 65 (3): 387–403.

Härtig, E., U. Schiek, K. U. Vollack, and W. G. Zumft. 1999. “Nitrate and Nitrite Control of Respiratory Nitrate Reduction in Denitrifying Pseudomonas Stutzeri by a Two-Component Regulatory System Homologous to NarXL of Escherichia Coli.” *Journal of Bacteriology* 181 (12): 3658–65.

Hirakawa, Hidetada, Kunihiko Nishino, Takahiro Hirata, and Akihito Yamaguchi. 2003. “Comprehensive Studies of Drug Resistance Mediated by Overexpression of Response Regulators of Two-Component Signal Transduction Systems in Escherichia Coli.” *Journal of Bacteriology* 185 (6): 1851–56.

Hirakawa, Hidetada, Kunihiko Nishino, Junko Yamada, Takahiro Hirata, and Akihito Yamaguchi. 2003. “Beta-Lactam Resistance Modulated by the Overexpression of Response Regulators of Two-Component Signal Transduction Systems in Escherichia Coli.” *The Journal of Antimicrobial Chemotherapy* 52 (4): 576–82.

Hobbs, M., E. S. Collie, P. D. Free, S. P. Livingston, and J. S. Mattick. 1993. “PilS and PilR, a Two-Component Transcriptional Regulatory System Controlling Expression of Type 4 Fimbriae in Pseudomonas Aeruginosa.” *Molecular Microbiology* 7 (5): 669–82.

Hong, Hee-Jeon, Matthew I. Hutchings, Mark J. Buttner, and Biotechnology and Biological Sciences Research Council, UK. 2008. “Vancomycin Resistance VanS/VanR Two-Component Systems.” *Advances in Experimental Medicine and Biology* 631: 200–213.

Hsieh, Yi-Ju, and Barry L. Wanner. 2010. “Global Regulation by the Seven-Component Pi Signaling System.” *Current Opinion in Microbiology* 13 (2): 198–203.

Hsu, Jye-Lin, Hsuan-Cheng Chen, Hwei-Ling Peng, and Hwan-You Chang. 2008. “Characterization of the Histidine-Containing Phosphotransfer Protein B-Mediated Multistep Phosphorelay System in Pseudomonas Aeruginosa PAO1.” *The Journal of Biological Chemistry* 283 (15): 9933–44.

Hurley, Bryan P., Andrew L. Goodman, Karen L. Mumy, Patrick Murphy, Stephen Lory, and Beth A. McCormick. 2010. “The Two-Component Sensor Response Regulator RoxS/RoxR Plays a Role in Pseudomonas Aeruginosa Interactions with Airway Epithelial Cells.” *Microbes and Infection / Institut Pasteur* 12 (3): 190–98.

Huynh, Tuanh Ngoc, Chris E. Noriega, and Valley Stewart. 2010. “Conserved

Mechanism for Sensor Phosphatase Control of Two-Component Signaling Revealed in the Nitrate Sensor NarX." *Proceedings of the National Academy of Sciences of the United States of America* 107 (49): 21140–45.
Jack, R., M. De Zamaroczy, and M. Merrick. 1999. "The Signal Transduction Protein GlnK Is Required for NifL-Dependent Nitrogen Control of Nif Gene Expression in Klebsiella Pneumoniae." *Journal of Bacteriology* 181 (4): 1156–62.
Jayol, Aurélie, Patrice Nordmann, Adrian Brink, and Laurent Poirel. 2015. "Heteroresistance to Colistin in Klebsiella Pneumoniae Associated with Alterations in the PhoPQ Regulatory System." *Antimicrobial Agents and Chemotherapy* 59 (5): 2780–84.
Jung, Kirsten, Florian Fabiani, Elisabeth Hoyer, and Jürgen Lassak. 2018. "Bacterial Transmembrane Signalling Systems and Their Engineering for Biosensing." *Open Biology* 8 (4). https://doi.org/10.1098/rsob.180023.
Kang, Katie N., Dustin R. Klein, Misha I. Kazi, François Guérin, Vincent Cattoir, Jennifer S. Brodbelt, and Joseph M. Boll. 2019. "Colistin Heteroresistance in Enterobacter Cloacae Is Regulated by PhoPQ-Dependent 4-Amino-4-Deoxy-L-Arabinose Addition to Lipid A." *Molecular Microbiology* 111 (6): 1604–16.
Kawakami, Takuro, Miho Kuroki, Masaharu Ishii, Yasuo Igarashi, and Hiroyuki Arai. 2010. "Differential Expression of Multiple Terminal Oxidases for Aerobic Respiration in Pseudomonas Aeruginosa." *Environmental Microbiology* 12 (6): 1399–1412.
Keller, R., M. Z. Pedroso, R. Ritchmann, and R. M. Silva. 1998. "Occurrence of Virulence-Associated Properties in Enterobacter Cloacae." *Infection and Immunity* 66 (2): 645–49.
Kilmury, Sara L. N., and Lori L. Burrows. 2018. "The Pseudomonas Aeruginosa PilSR Two-Component System Regulates Both Twitching and Swimming Motilities." *mBio* 9 (4). https://doi.org/10.1128/mBio.01310-18.
Kim, Jung Wook, Hyun-Kyung Kim, Gi Su Kang, Il-Hwan Kim, Hwa Su Kim, Yeong Seon Lee, and Jae Il Yoo. 2016. "The SAV1322 Gene from Staphylococcus Aureus: Genomic and Proteomic Approaches to Identification and Characterization of Gene Function." *BMC Microbiology* 16 (1): 206.
Kim, So Yeon, Hyeon Jin Choi, and Kwan Soo Ko. 2014. "Differential Expression of Two-Component Systems, pmrAB and phoPQ, with Different Growth Phases of Klebsiella Pneumoniae in the Presence or Absence of Colistin." *Current Microbiology* 69 (1): 37–41.
Kinkel, Traci L., Christelle M. Roux, Paul M. Dunman, and Ferric C. Fang. 2013. "The Staphylococcus Aureus SrrAB Two-Component System Promotes Resistance to Nitrosative Stress and Hypoxia." *mBio* 4 (6): e00696–13.
Knopp, Michael, and Dan I. Andersson. 2015. "Amelioration of the Fitness Costs of Antibiotic Resistance Due To Reduced Outer Membrane Permeability by Upregulation of Alternative Porins." *Molecular Biology and Evolution* 32 (12): 3252–63.
Kraxenberger, Tobias, Luitpold Fried, Stefan Behr, and Kirsten Jung. 2012. "First Insights into the Unexplored Two-Component System YehU/YehT in Escherichia Coli." *Journal of Bacteriology* 194 (16): 4272–84.

Kremer, Anita N., and Harald Hoffmann. 2012. “Subtractive Hybridization Yields a Silver Resistance Determinant Unique to Nosocomial Pathogens in the Enterobacter Cloacae Complex.” *Journal of Clinical Microbiology* 50 (10): 3249–57.

Kumar, Vinod, Peng Sun, Jessica Vamathevan, Yong Li, Karen Ingraham, Leslie Palmer, Jianzhong Huang, and James R. Brown. 2011. “Comparative Genomics of Klebsiella Pneumoniae Strains with Different Antibiotic Resistance Profiles.” *Antimicrobial Agents and Chemotherapy* 55 (9): 4267–76.

Kurabayashi, Kumiko, Yuko Hirakawa, Koichi Tanimoto, Haruyoshi Tomita, and Hidetada Hirakawa. 2014. “Role of the CpxAR Two-Component Signal Transduction System in Control of Fosfomycin Resistance and Carbon Substrate Uptake.” *Journal of Bacteriology* 196 (2): 248–56.

Kuroda, Makoto, Hiroko Kuroda, Taku Oshima, Fumihiko Takeuchi, Hirotada Mori, and Keiichi Hiramatsu. 2003. “Two-Component System VraSR Positively Modulates the Regulation of Cell-Wall Biosynthesis Pathway in Staphylococcus Aureus.” *Molecular Microbiology* 49 (3): 807–21.

Kyriakidis, Dimitrios A., and Ekaterini Tiligada. 2009. “Signal Transduction and Adaptive Regulation through Bacterial Two-Component Systems: The Escherichia Coli AtoSC Paradigm.” *Amino Acids* 37 (3): 443–58.

Lai, Yi-Chyi, Ging-Ting Lin, Shu-Li Yang, Hwan-You Chang, and Hwei-Ling Peng. 2003. “Identification and Characterization of KvgAS, a Two-Component System in Klebsiella Pneumoniae CG43.” *FEMS Microbiology Letters* 218 (1): 121–26.

Landry, Brian P., Rohan Palanki, Nikola Dyulgyarov, Lucas A. Hartsough, and Jeffrey J. Tabor. 2018. “Phosphatase Activity Tunes Two-Component System Sensor Detection Threshold.” *Nature Communications* 9 (1): 1433.

Lau, Calvin Ho-Fung, Sebastien Fraud, Marcus Jones, Scott N. Peterson, and Keith Poole. 2013. “Mutational Activation of the AmgRS Two-Component System in Aminoglycoside-Resistant Pseudomonas Aeruginosa.” *Antimicrobial Agents and Chemotherapy* 57 (5): 2243–51.

Lautenbach, E., J. B. Patel, W. B. Bilker, P. H. Edelstein, and N. O. Fishman. 2001. “Extended-Spectrum Beta-Lactamase-Producing Escherichia Coli and Klebsiella Pneumoniae: Risk Factors for Infection and Impact of Resistance on Outcomes.” *Clinical Infectious Diseases: An Official Publication of the Infectious Diseases Society of America* 32 (8): 1162–71.

Lazar Adler, Natalie R., Elizabeth M. Allwood, Deanna Deveson Lucas, Paul Harrison, Stephen Watts, Alexandra Dimitropoulos, Puthayalai Treerat, et al. 2016. “Perturbation of the Two-Component Signal Transduction System, BprRS, Results in Attenuated Virulence and Motility Defects in Burkholderia Pseudomallei.” *BMC Genomics* 17 (May): 331.

Leblanc, Shannon K. D., Christopher W. Oates, and Tracy L. Raivio. 2011. “Characterization of the Induction and Cellular Role of the BaeSR Two-Component Envelope Stress Response of Escherichia Coli.” *Journal of Bacteriology* 193 (13): 3367–75.

Leday, Temekka V., Kathryn M. Gold, Traci L. Kinkel, Samantha A. Roberts, June R. Scott, and Kevin S. McIver. 2008. “TrxR, a New CovR-Repressed Response Regulator That Activates the Mga Virulence Regulon in Group A Streptococcus.”

*Infection and Immunity* 76 (10): 4659–68.

Lesne, E., E-M Krammer, E. Dupre, C. Locht, M. F. Lensink, R. Antoine, and F. Jacob-Dubuisson. 2016. “Balance between Coiled-Coil Stability and Dynamics Regulates Activity of BvgS Sensor Kinase in Bordetella.” *mBio* 7 (2): e02089.

Li, Congran, Xiangyu Yao, and Chung-Dar Lu. 2010. “Regulation of the dauBAR Operon and Characterization of D-Amino Acid Dehydrogenase DauA in Arginine and Lysine Catabolism of Pseudomonas Aeruginosa PAO1.” *Microbiology* 156 (Pt 1): 60–71.

Lin, Ching-Ting, and Hwei-Ling Peng. 2006. “Regulation of the Homologous Two-Component Systems KvgAS and KvhAS in Klebsiella Pneumoniae CG43.” *Journal of Biochemistry* 140 (5): 639–48.

Lin, Ming-Feng, Yun-You Lin, Hui-Wen Yeh, and Chung-Yu Lan. 2014. “Role of the BaeSR Two-Component System in the Regulation of Acinetobacter Baumannii adeAB Genes and Its Correlation with Tigecycline Susceptibility.” *BMC Microbiology* 14 (May): 119.

Lin, Tien-Huang, Yeh Chen, Jong-Tar Kuo, Yi-Chyi Lai, Chien-Chen Wu, Chun-Fa Huang, and Ching-Ting Lin. 2018. “Phosphorylated OmpR Is Required for Type 3 Fimbriae Expression in Under Hypertonic Conditions.” *Frontiers in Microbiology* 9 (October): 2405.

Li, Wei, and Chung-Dar Lu. 2007. “Regulation of Carbon and Nitrogen Utilization by CbrAB and NtrBC Two-Component Systems in Pseudomonas Aeruginosa.” *Journal of Bacteriology* 189 (15): 5413–20.

Macielag, M. J., J. P. Demers, S. A. Fraga-Spano, D. J. Hlasta, S. G. Johnson, R. M. Kanojia, R. K. Russell, et al. 1998. “Substituted Salicylanilides as Inhibitors of Two-Component Regulatory Systems in Bacteria.” *Journal of Medicinal Chemistry* 41 (16): 2939–45.

Majewski, Piotr, Piotr Wieczorek, Dominika Ojdana, Anna Sieńko, Oksana Kowalczuk, Paweł Sacha, Jacek Nikliński, and Elżbieta Tryniszewska. 2016. “Altered Outer Membrane Transcriptome Balance with AmpC Overexpression in Carbapenem-Resistant.” *Frontiers in Microbiology* 7 (December): 2054.

Makarova, Olga, Paul Johnston, Alexandro Rodriguez-Rojas, Baydaa El Shazely, Javier Moreno Morales, and Jens Rolff. 2018. “Genomics of Experimental Adaptation of Staphylococcus Aureus to a Natural Combination of Insect Antimicrobial Peptides.” *Scientific Reports* 8 (1): 15359.

Ma, Siya, Siwei Luo, L. I. Wu, Zhi Liang, and Jia-Rui Wu. 2017. “Re-Engineering the Two-Component Systems as Light-Regulated in.” *Journal of Biosciences* 42 (4): 565–73.

Masuda, Nobuhisa, and George M. Church. 2002. “Escherichia Coli Gene Expression Responsive to Levels of the Response Regulator EvgA.” *Journal of Bacteriology* 184 (22): 6225–34.

Mensa, Bruk, Gabriella L. Howell, Richard Scott, and William F. DeGrado. 2014. “Comparative Mechanistic Studies of Brilacidin, Daptomycin, and the Antimicrobial Peptide LL16.” *Antimicrobial Agents and Chemotherapy* 58 (9): 5136–45.

Mern, Demissew S., Seung-Wook Ha, Viola Khodaverdi, Nicole Gliese, and Helmut Görisch. 2010. “A Complex Regulatory Network Controls Aerobic Ethanol Oxidation in Pseudomonas Aeruginosa: Indication of Four Levels of Sensor Kinases and Response Regulators.” *Microbiology* 156 (Pt 5): 1505–16.

Mikkelsen, Helga, Geneviève Ball, Caroline Giraud, and Alain Filloux. 2009. “Expression of Pseudomonas Aeruginosa CupD Fimbrial Genes Is Antagonistically Controlled by RcsB and the EAL-Containing PvrR Response Regulators.” *PloS One* 4 (6): e6018.

Mikkelsen, Helga, Kailyn Hui, Nicolas Barraud, and Alain Filloux. 2013. “The Pathogenicity Island Encoded PvrSR/RcsCB Regulatory Network Controls Biofilm Formation and Dispersal in Pseudomonas Aeruginosa PA14.” *Molecular Microbiology* 89 (3): 450–63.

Minagawa, Shu, Hiroshi Ogasawara, Akinori Kato, Kaneyoshi Yamamoto, Yoko Eguchi, Taku Oshima, Hirotada Mori, Akira Ishihama, and Ryutaro Utsumi. 2003. “Identification and Molecular Characterization of the Mg2+ Stimulon of Escherichia Coli.” *Journal of Bacteriology* 185 (13): 3696–3702.

Miyake, Yukari, Tatsuya Inaba, Hiroki Watanabe, Jun Teramoto, Kaneyoshi Yamamoto, and Akira Ishihama. 2019. “Regulatory Roles of Pyruvate-Sensing Two-Component System PyrSR (YpdAB) in Escherichia Coli K-12.” *FEMS Microbiology Letters* 366 (2). https://doi.org/10.1093/femsle/fnz009.

Miyashiro, Tim, and Mark Goulian. 2007. “Stimulus-Dependent Differential Regulation in the Escherichia Coli PhoQ PhoP System.” *Proceedings of the National Academy of Sciences of the United States of America* 104 (41): 16305–10.

Muller, Cédric, Patrick Plésiat, and Katy Jeannot. 2011. “A Two-Component Regulatory System Interconnects Resistance to Polymyxins, Aminoglycosides, Fluoroquinolones, and β-Lactams in Pseudomonas Aeruginosa.” *Antimicrobial Agents and Chemotherapy* 55 (3): 1211–21.

Munson, G. P., D. L. Lam, F. W. Outten, and T. V. O’Halloran. 2000. “Identification of a Copper-Responsive Two-Component System on the Chromosome of Escherichia Coli K-12.” *Journal of Bacteriology* 182 (20): 5864–71.

Muzamal, Uzma, Daniel Gomez, Fenika Kapadia, and Dasantila Golemi-Kotra. 2014. “Diversity of Two-Component Systems: Insights into the Signal Transduction Mechanism by the Staphylococcus Aureus Two-Component System GraSR.” *F1000Research* 3 (October): 252.

Ninfa, A. J., and B. Magasanik. 1986. “Covalent Modification of the glnG Product, NRI, by the glnL Product, NRII, Regulates the Transcription of the glnALG Operon in Escherichia Coli.” *Proceedings of the National Academy of Sciences of the United States of America* 83 (16): 5909–13.

Nishino, Kunihiko, Takeshi Honda, and Akihito Yamaguchi. 2005. “Genome-Wide Analyses of Escherichia Coli Gene Expression Responsive to the BaeSR Two-Component Regulatory System.” *Journal of Bacteriology* 187 (5): 1763–72.

Niu, S., S. Q. Jiang, and J. Hong. 1995. “Salmonella Typhimurium pgtB Mutants Conferring Constitutive Expression of Phosphoglycerate Transporter pgtP Independent of pgtC.” *Journal of Bacteriology* 177 (15): 4297–4302.

Nixon, B. T., C. W. Ronson, and F. M. Ausubel. 1986. “Two-Component Regulatory Systems Responsive to Environmental Stimuli Share Strongly Conserved Domains with the Nitrogen Assimilation Regulatory Genes ntrB and ntrC.” *Proceedings of the*

*National Academy of Sciences of the United States of America* 83 (20): 7850–54.
Novick, Richard P., and Dunrong Jiang. 2003. “The Staphylococcal saeRS System Coordinates Environmental Signals with Agr Quorum Sensing.” *Microbiology* 149 (Pt 10): 2709–17.
Novick, R. P., S. J. Projan, J. Kornblum, H. F. Ross, G. Ji, B. Kreiswirth, F. Vandenesch, and S. Moghazeh. 1995. “The Agr P2 Operon: An Autocatalytic Sensory Transduction System in Staphylococcus Aureus.” *Molecular & General Genetics: MGG* 248 (4): 446–58.
Ogasawara, Hiroshi, Shota Shinohara, Kaneyoshi Yamamoto, and Akira Ishihama. 2012. “Novel Regulation Targets of the Metal-Response BasS-BasR Two-Component System of Escherichia Coli.” *Microbiology* 158 (Pt 6): 1482–92.
Okkotsu, Yuta, Alexander S. Little, and Michael J. Schurr. 2014. “The Pseudomonas Aeruginosa AlgZR Two-Component System Coordinates Multiple Phenotypes.” *Frontiers in Cellular and Infection Microbiology* 4 (June): 82.
Panesso, Diana, Jinnethe Reyes, Elizabeth P. Gaston, Morgan Deal, Alejandra Londoño, Masayuki Nigo, Jose M. Munita, et al. 2015. “Deletion of liaR Reverses Daptomycin Resistance in Enterococcus Faecium Independent of the Genetic Background.” *Antimicrobial Agents and Chemotherapy* 59 (12): 7327–34.
Petit-Härtlein, Isabelle, Kevin Rome, Eve de Rosny, Florian Molton, Carole Duboc, Erwan Gueguen, Agnès Rodrigue, and Jacques Covès. 2015. “Biophysical and Physiological Characterization of ZraP from Escherichia Coli, the Periplasmic Accessory Protein of the Atypical ZraSR Two-Component System.” *Biochemical Journal* 472 (2): 205–16.
Petrova, Olga E., and Karin Sauer. 2010. “The Novel Two-Component Regulatory System BfiSR Regulates Biofilm Development by Controlling the Small RNA rsmZ through CafA.” *Journal of Bacteriology* 192 (20): 5275–88.
Porcheron, Gaëlle, Amélie Garénaux, Julie Proulx, Mourad Sabri, and Charles M. Dozois. 2013. “Iron, Copper, Zinc, and Manganese Transport and Regulation in Pathogenic Enterobacteria: Correlations between Strains, Site of Infection and the Relative Importance of the Different Metal Transport Systems for Virulence.” *Frontiers in Cellular and Infection Microbiology* 3 (December): 90.
Pragman, Alexa A., Jeremy M. Yarwood, Timothy J. Tripp, and Patrick M. Schlievert. 2004. “Characterization of Virulence Factor Regulation by SrrAB, a Two-Component System in Staphylococcus Aureus.” *Journal of Bacteriology* 186 (8): 2430–38.
Pritchett, Christopher L., Alexander S. Little, Yuta Okkotsu, Anders Frisk, William L. Cody, Christopher R. Covey, and Michael J. Schurr. 2015. “Expression Analysis of the Pseudomonas Aeruginosa AlgZR Two-Component Regulatory System.” *Journal of Bacteriology* 197 (4): 736–48.
Radin, Jana N., Jessica L. Kelliher, Paola K. Párraga Solórzano, and Thomas E. Kehl-Fie. 2016. “The Two-Component System ArlRS and Alterations in Metabolism Enable Staphylococcus Aureus to Resist Calprotectin-Induced Manganese Starvation.” *PLoS Pathogens* 12 (11): e1006040.
Raivio, Tracy L. 2014. “Everything Old Is New Again: An Update on Current Research on the Cpx Envelope Stress Response.” *Biochimica et Biophysica Acta* 1843 (8):

1529–41.
Ramos, Pablo Ivan Pereira, Márlon Grégori Flores Custódio, Guadalupe Del Rosario Quispe Saji, Thiago Cardoso, Gisele Lucchetti da Silva, Graziela Braun, Willames M. B. S. Martins, et al. 2016. “The Polymyxin B-Induced Transcriptomic Response of a Clinical, Multidrug-Resistant Klebsiella Pneumoniae Involves Multiple Regulatory Elements and Intracellular Targets.” *BMC Genomics* 17 (Suppl 8): 737.
Ravikumar, Sambandam, Van Dung Pham, Seung Hwan Lee, Ik-Keun Yoo, and Soon Ho Hong. 2012. “Modification of CusSR Bacterial Two-Component Systems by the Introduction of an Inducible Positive Feedback Loop.” *Journal of Industrial Microbiology & Biotechnology* 39 (6): 861–68.
Ren, Yan, Yi Ren, Zhemin Zhou, Xi Guo, Yayue Li, Lu Feng, and Lei Wang. 2010. “Complete Genome Sequence of Enterobacter Cloacae Subsp. Cloacae Type Strain ATCC 13047.” *Journal of Bacteriology* 192 (9): 2463–64.
Richmond, Grace E., Laura P. Evans, Michele J. Anderson, Matthew E. Wand, Laura C. Bonney, Alasdair Ivens, Kim Lee Chua, et al. 2016. “The Acinetobacter Baumannii Two-Component System AdeRS Regulates Genes Required for Multidrug Efflux, Biofilm Formation, and Virulence in a Strain-Specific Manner.” *mBio* 7 (2): e00430–16.
Roberts, Daniel P., Scott M. Lohrke, Laurie McKenna, Dilip K. Lakshman, Hyesuk Kong, and John Lydon. 2011. “Mutation of a degS Homologue in Enterobacter Cloacae Decreases Colonization and Biological Control of Damping-off on Cucumber.” *Phytopathology* 101 (2): 271–80.
Rome, Kévin, Céline Borde, Raleb Taher, Julien Cayron, Christian Lesterlin, Erwan Gueguen, Eve De Rosny, and Agnès Rodrigue. 2018. “The Two-Component System ZraPSR Is a Novel ESR That Contributes to Intrinsic Antibiotic Tolerance in Escherichia Coli.” *Journal of Molecular Biology* 430 (24): 4971–85.
Samir, Reham, Salma H. Hussein, Noha M. Elhosseiny, Marwa S. Khattab, Alaa E. Shawky, and Ahmed S. Attia. 2016. “Adaptation to Potassium-Limitation Is Essential for Acinetobacter Baumannii Pneumonia Pathogenesis.” *The Journal of Infectious Diseases* 214 (12): 2006–13.
Santos-Beneit, Fernando. 2015. “The Pho Regulon: A Huge Regulatory Network in Bacteria.” *Frontiers in Microbiology* 6 (April): 402.
Scheu, P. D., J. Witan, M. Rauschmeier, S. Graf, Y-F Liao, A. Ebert-Jung, T. Basché, W. Erker, and G. Unden. 2012. “CitA/CitB Two-Component System Regulating Citrate Fermentation in Escherichia Coli and Its Relation to the DcuS/DcuR System in Vivo.” *Journal of Bacteriology* 194 (3): 636–45.
Schlag, Steffen, Stephan Fuchs, Christiane Nerz, Rosmarie Gaupp, Susanne Engelmann, Manuel Liebeke, Michael Lalk, Michael Hecker, and Friedrich Götz. 2008. “Characterization of the Oxygen-Responsive NreABC Regulon of Staphylococcus Aureus.” *Journal of Bacteriology* 190 (23): 7847–58.
Schmidl, Sebastian R., Felix Ekness, Katri Sofjan, Kristina N-M Daeffler, Kathryn R. Brink, Brian P. Landry, Karl P. Gerhardt, Nikola Dyulgyarov, Ravi U. Sheth, and Jeffrey J. Tabor. 2019. “Rewiring Bacterial Two-Component Systems by Modular DNA-Binding Domain Swapping.” *Nature Chemical Biology* 15 (7): 690–98.
Shankar, Manoharan, Saswat S. Mohapatra, Saswati Biswas, and Indranil Biswas.

2015. “Gene Regulation by the LiaSR Two-Component System in Streptococcus Mutans.” *PloS One* 10 (5): e0128083.
Sharma-Kuinkel, Batu K., Ethan E. Mann, Jong-Sam Ahn, Lisa J. Kuechenmeister, Paul M. Dunman, and Kenneth W. Bayles. 2009. “The Staphylococcus Aureus LytSR Two-Component Regulatory System Affects Biofilm Formation.” *Journal of Bacteriology* 191 (15): 4767–75.
Singh, Kamna, Dilani B. Senadheera, and Dennis G. Cvitkovitch. 2014. “An Intimate Link: Two-Component Signal Transduction Systems and Metal Transport Systems in Bacteria.” *Future Microbiology* 9 (11): 1283–93.
Sobran, M. Ashley, and Peggy A. Cotter. 2019. “The BvgS PAS Domain, an Independent Sensory Perception Module in the BvgAS Phosphorelay.” *Journal of Bacteriology* 201 (17). https://doi.org/10.1128/JB.00286-19.
Sonawane, Avinash M., Birendra Singh, and Klaus-Heinrich Röhm. 2006. “The AauR-AauS Two-Component System Regulates Uptake and Metabolism of Acidic Amino Acids in Pseudomonas Putida.” *Applied and Environmental Microbiology* 72 (10): 6569–77.
Srinivasan, Vijaya Bharathi, Vasanth Vaidyanathan, Amitabha Mondal, and Govindan Rajamohan. 2012. “Role of the Two Component Signal Transduction System CpxAR in Conferring Cefepime and Chloramphenicol Resistance in Klebsiella Pneumoniae NTUH-K2044.” *PloS One* 7 (4): e33777.
Srinivasan, Vijaya Bharathi, Manjunath Venkataramaiah, Amitabha Mondal, Vasanth Vaidyanathan, Tanvi Govil, and Govindan Rajamohan. 2012. “Functional Characterization of a Novel Outer Membrane Porin KpnO, Regulated by PhoBR Two-Component System in Klebsiella Pneumoniae NTUH-K2044.” *PloS One* 7 (7): e41505.
Staehlin, Benjamin M., John G. Gibbons, Antonis Rokas, Thomas V. O’Halloran, and Jason C. Slot. 2016. “Evolution of a Heavy Metal Homeostasis/Resistance Island Reflects Increasing Copper Stress in Enterobacteria.” *Genome Biology and Evolution* 8 (3): 811–26.
Stauff, Devin L., Victor J. Torres, and Eric P. Skaar. 2007. “Signaling and DNA-Binding Activities of the Staphylococcus Aureus HssR-HssS Two-Component System Required for Heme Sensing.” *The Journal of Biological Chemistry* 282 (36): 26111–21.
Su, Kewen, Xipeng Zhou, Mei Luo, Xuan Xu, Pin Liu, Xuan Li, Jian Xue, et al. 2018. “Genome-Wide Identification of Genes Regulated by RcsA, RcsB, and RcsAB Phosphorelay Regulators in Klebsiella Pneumoniae NTUH-K2044.” *Microbial Pathogenesis* 123 (October): 36–41.
Sun, Junsong, Li Zheng, Christina Landwehr, Junshu Yang, and Yinduo Ji. 2005. “Identification of a Novel Essential Two-Component Signal Transduction System, YhcSR, in Staphylococcus Aureus.” *Journal of Bacteriology* 187 (22): 7876–80.
Sun, Song, Aurel Negrea, Mikael Rhen, and Dan I. Andersson. 2009. “Genetic Analysis of Colistin Resistance in Salmonella Enterica Serovar Typhimurium.” *Antimicrobial Agents and Chemotherapy* 53 (6): 2298–2305.
Szklarczyk, Damian, Annika L. Gable, David Lyon, Alexander Junge, Stefan Wyder, Jaime Huerta-Cepas, Milan Simonovic, et al. 2019. “STRING v11: Protein-Protein

Association Networks with Increased Coverage, Supporting Functional Discovery in Genome-Wide Experimental Datasets." *Nucleic Acids Research* 47 (D1): D607–13.
Tatke, Gorakh, Hansi Kumari, Eugenia Silva-Herzog, Lourdes Ramirez, and Kalai Mathee. 2015. "Pseudomonas Aeruginosa MifS-MifR Two-Component System Is Specific for α-Ketoglutarate Utilization." *PloS One* 10 (6): e0129629.
Taylor, Patrick K., Li Zhang, and Thien-Fah Mah. 2019. "Loss of the Two-Component System TctD-TctE in Affects Biofilm Formation and Aminoglycoside Susceptibility in Response to Citric Acid." *mSphere* 4 (2). https://doi.org/10.1128/mSphere.00102-19.
Theodorou, Marina C., Evaggelos C. Theodorou, and Dimitrios A. Kyriakidis. 2012. "Involvement of AtoSC Two-Component System in Escherichia Coli Flagellar Regulon." *Amino Acids* 43 (2): 833–44.
Tipton, Kyle A., and Philip N. Rather. 2017. "An Two-Component System Ortholog Regulates Phase Variation, Osmotic Tolerance, Motility, and Virulence in Acinetobacter Baumannii Strain AB5075." *Journal of Bacteriology* 199 (3). https://doi.org/10.1128/JB.00705-16.
Tiwari, Sandeep, Syed B. Jamal, Syed S. Hassan, Paulo V. S. D. Carvalho, Sintia Almeida, Debmalya Barh, Preetam Ghosh, Artur Silva, Thiago L. P. Castro, and Vasco Azevedo. 2017. "Two-Component Signal Transduction Systems of Pathogenic Bacteria As Targets for Antimicrobial Therapy: An Overview." *Frontiers in Microbiology* 8 (October): 1878.
Toro-Roman, Alejandro, Ti Wu, and Ann M. Stock. 2005. "A Common Dimerization Interface in Bacterial Response Regulators KdpE and TorR." *Protein Science: A Publication of the Protein Society* 14 (12): 3077–88.
Torres, Victor J., Devin L. Stauff, Gleb Pishchany, Jelena S. Bezbradica, Laura E. Gordy, Juan Iturregui, Kelsi L. Anderson, Paul M. Dunman, Sebastian Joyce, and Eric P. Skaar. 2007. "A Staphylococcus Aureus Regulatory System That Responds to Host Heme and Modulates Virulence." *Cell Host & Microbe* 1 (2): 109–19.
Tran, Truc T., Diana Panesso, Hongyu Gao, Jung H. Roh, Jose M. Munita, Jinnethe Reyes, Lorena Diaz, et al. 2013. "Whole-Genome Analysis of a Daptomycin-Susceptible Enterococcus Faecium Strain and Its Daptomycin-Resistant Variant Arising during Therapy." *Antimicrobial Agents and Chemotherapy* 57 (1): 261–68.
Trastoy, R., T. Manso, L. Fernández-García, L. Blasco, A. Ambroa, M. L. Pérez Del Molino, G. Bou, R. García-Contreras, T. K. Wood, and M. Tomás. 2018. "Mechanisms of Bacterial Tolerance and Persistence in the Gastrointestinal and Respiratory Environments." *Clinical Microbiology Reviews* 31 (4). https://doi.org/10.1128/CMR.00023-18.
Udaondo, Zulema, Juan-Luis Ramos, Ana Segura, Tino Krell, and Abdelali Daddaoua. 2018. "Regulation of Carbohydrate Degradation Pathways in Pseudomonas Involves a Versatile Set of Transcriptional Regulators." *Microbial Biotechnology* 11 (3): 442–54.
Unden, G., S. Wörner, and C. Monzel. 2016. "Cooperation of Secondary Transporters and Sensor Kinases in Transmembrane Signalling: The DctA/DcuS and DcuB/DcuS Sensor Complexes of Escherichia Coli." *Advances in Microbial*

*Physiology* 68 (March): 139–67.
Urano, Hiroyuki, Myu Yoshida, Ayano Ogawa, Kaneyoshi Yamamoto, Akira Ishihama, and Hiroshi Ogasawara. 2017. "Cross-Regulation between Two Common Ancestral Response Regulators, HprR and CusR, in Escherichia Coli." *Microbiology* 163 (2): 243–52.
Verhamme, Daniël T., Pieter W. Postma, Wim Crielaard, and Klaas J. Hellingwerf. 2002. "Cooperativity in Signal Transfer through the Uhp System of Escherichia Coli." *Journal of Bacteriology* 184 (15): 4205–10.
Villanueva, Maite, Begoña García, Jaione Valle, Beatriz Rapún, Igor Ruiz de Los Mozos, Cristina Solano, Miguel Martí, José R. Penadés, Alejandro Toledo-Arana, and Iñigo Lasa. 2018. "Sensory Deprivation in Staphylococcus Aureus." *Nature Communications* 9 (1): 523.
Walker, Jennifer N., Heidi A. Crosby, Adam R. Spaulding, Wilmara Salgado-Pabón, Cheryl L. Malone, Carolyn B. Rosenthal, Patrick M. Schlievert, Jeffrey M. Boyd, and Alexander R. Horswill. 2013. "The Staphylococcus Aureus ArlRS Two-Component System Is a Novel Regulator of Agglutination and Pathogenesis." *PLoS Pathogens* 9 (12): e1003819.
Wanner, B. L. 1992. "Is Cross Regulation by Phosphorylation of Two-Component Response Regulator Proteins Important in Bacteria?" *Journal of Bacteriology* 174 (7): 2053–58.
Weigel, W. A., and D. R. Demuth. 2016. "QseBC, a Two-Component Bacterial Adrenergic Receptor and Global Regulator of Virulence in Enterobacteriaceae and Pasteurellaceae." *Molecular Oral Microbiology* 31 (5): 379–97.
Wen, Yurong, Zhenlin Ouyang, Yue Yu, Xiaorong Zhou, Yingmei Pei, Bart Devreese, Paul G. Higgins, and Fang Zheng. 2017. "Mechanistic Insight into How Multidrug Resistant Acinetobacter Baumannii Response Regulator AdeR Recognizes an Intercistronic Region." *Nucleic Acids Research* 45 (16): 9773–87.
Weston, L. A., and R. J. Kadner. 1988. "Role of Uhp Genes in Expression of the Escherichia Coli Sugar-Phosphate Transport System." *Journal of Bacteriology* 170 (8): 3375–83.
Willett, Jonathan W., and Sean Crosson. 2017. "Atypical Modes of Bacterial Histidine Kinase Signaling." *Molecular Microbiology* 103 (2): 197–202.
Wu, Fei, Yuanyuan Ying, Min Yin, Yi Jiang, Chongyang Wu, Changrui Qian, Qianqian Chen, et al. 2019. "Molecular Characterization of a Multidrug-Resistant Strain R46 Isolated from a Rabbit." *International Journal of Genomics and Proteomics* 2019 (August): 5459190.
Xue, Ting, Yibo You, De Hong, Haipeng Sun, and Baolin Sun. 2011. "The Staphylococcus Aureus KdpDE Two-Component System Couples Extracellular K+ Sensing and Agr Signaling to Infection Programming." *Infection and Immunity* 79 (6): 2154–67.
Yamamoto, Kaneyoshi, Kiyo Hirao, Taku Oshima, Hirofumi Aiba, Ryutaro Utsumi, and Akira Ishihama. 2005. "Functional Characterization in Vitro of All Two-Component Signal Transduction Systems from Escherichia Coli." *The Journal of Biological Chemistry* 280 (2): 1448–56.
Yang, Soo-Jin, Arnold S. Bayer, Nagendra N. Mishra, Michael Meehl, Nagender Ledala,

Michael R. Yeaman, Yan Q. Xiong, and Ambrose L. Cheung. 2012. “The Staphylococcus Aureus Two-Component Regulatory System, GraRS, Senses and Confers Resistance to Selected Cationic Antimicrobial Peptides.” *Infection and Immunity* 80 (1): 74–81.
Yan, Meiying, Jeffrey W. Hall, Junshu Yang, and Yinduo Ji. 2012. “The Essential yhcSR Two-Component Signal Transduction System Directly Regulates the Lac and opuCABCD Operons of Staphylococcus Aureus.” *PloS One* 7 (11): e50608.
Yin, Shaohui, Robert S. Daum, and Susan Boyle-Vavra. 2006. “VraSR Two-Component Regulatory System and Its Role in Induction of pbp2 and vraSR Expression by Cell Wall Antimicrobials in Staphylococcus Aureus.” *Antimicrobial Agents and Chemotherapy* 50 (1): 336–43.
Yuan, Jie, Buyun Wei, Miaomiao Shi, and Haichun Gao. 2011. “Functional Assessment of EnvZ/OmpR Two-Component System in Shewanella Oneidensis.” *PloS One* 6 (8): e23701.
Zahid, Nageena, Soumble Zulfiqar, and Abdul Rauf Shakoori. 2012. “Functional Analysis of Cus Operon Promoter of Klebsiella Pneumoniae Using E. Coli lacZ Assay.” *Gene* 495 (1): 81–88.
Zamorano, Laura, Bartolomé Moyà, Carlos Juan, Xavier Mulet, Jesús Blázquez, and Antonio Oliver. 2014. “The Pseudomonas Aeruginosa CreBC Two-Component System Plays a Major Role in the Response to β-Lactams, Fitness, Biofilm Growth, and Global Regulation.” *Antimicrobial Agents and Chemotherapy* 58 (9): 5084–95.
Zhao, Liping, Ting Xue, Fei Shang, Haipeng Sun, and Baolin Sun. 2010. “Staphylococcus Aureus AI-2 Quorum Sensing Associates with the KdpDE Two-Component System to Regulate Capsular Polysaccharide Synthesis and Virulence.” *Infection and Immunity* 78 (8): 3506–15.
Zhou, Lu, Xiang-He Lei, Barry R. Bochner, and Barry L. Wanner. 2003. “Phenotype Microarray Analysis of Escherichia Coli K-12 Mutants with Deletions of All Two-Component Systems.” *Journal of Bacteriology* 185 (16): 4956–72.
Zientz, E., J. Bongaerts, and G. Unden. 1998. “Fumarate Regulation of Gene Expression in Escherichia Coli by the DcuSR (dcuSR Genes) Two-Component Regulatory System.” *Journal of Bacteriology* 180 (20): 5421–25.
Zschiedrich, Christopher P., Victoria Keidel, and Hendrik Szurmant. 2016. “Molecular Mechanisms of Two-Component Signal Transduction.” *Journal of Molecular Biology* 428 (19): 3752–75.
